# Pre-stimulus cortical state predicts context-dependent expression of learned distractor suppression

**DOI:** 10.64898/2026.04.22.720064

**Authors:** Siyi Chen, Fredrik Allenmark, Hao Yu, Hermann J. Müller, Zhuanghua Shi

## Abstract

Suppressing visual distraction depends on learned attentional settings and the neural state preceding stimulus onset. We reanalyzed an fMRI visual-search dataset (N = 34) to test whether pre-stimulus global signal (GS), a broad state-sensitive BOLD measure, predicts subsequent performance and cortical responsivity across different learned distractor-suppression contexts. Participants searched for an orientation target while ignoring same-dimension orientation or different-dimension color distractors. Spatial regularity applied to the same-dimension distractor in one group and the different-dimension distractor in the other. Same-dimension distractors produced strong behavioral interference and recruited Dorsal Attention and Salience/Ventral Attention networks, whereas different-dimension distractors produced little interference. Spatial learning was also clearer when regularity applied to the target-relevant same dimension. Within these learning contexts, pre-stimulus GS predicted response speed in opposite directions across groups: higher GS preceded slower responses in the same-dimension spatial-bias group but faster responses in the different-dimension spatial-bias group. And pre-stimulus Default Mode Network activity showed a convergent context-dependent pattern. Higher pre-stimulus GS preceded reduced activity across Dorsal Attention, Salience/Ventral Attention, Control, Early Visual, and Default Mode regions. Although this neural attenuation was broadly expressed across conditions and groups, its association with the GS–RT relationship differed between learning contexts, most clearly on different-dimension trials. Together, these findings indicate that pre-stimulus GS reflects a broad modulation of subsequent cortical responsivity, whereas its behavioral consequences—and their relationship to regional neural attenuation—depend on the attentional-control context established by distractor dimension and spatial regularity.

## Introduction

Search begins with an aim. Whether we look for a friend in a crowd, a hazard in traffic, or a target in a display, attention must give priority to what matters and suppress what does not. The scene itself does not settle this competition. Current goals, selection history, and stimulus-driven priority jointly determine which objects guide selection and which are suppressed (Braver, 2012; Gaspelin & Luck, 2018; Wang & Theeuwes, 2018; Liesefeld et al., 2024). Ongoing brain state may then determine how strongly those settings govern the next act of selection. The same distractor may be suppressed efficiently in one moment and slow the same observer in the next, even when the external input is unchanged. The present study asks whether pre-stimulus cortical state modulates the expression of learned distractor suppression.

Visual search provides a strong test of this question because the functional architecture of distractor interference is well characterized (Found and Müller, 1996; Geyer et al., 2008; Kumada, 1999; Müller et al., 2009; Sauter et al., 2018). Salient distractors interfere most strongly when they share the target’s defining feature dimension. An orientation distractor, for example, is especially costly during orientation-guided search, whereas an equally salient color distractor can often be handled more efficiently when color is irrelevant. The dimension-weighting account explains this pattern by proposing that attentional weight is increased for the task-relevant dimension while it is reduced for irrelevant dimensions (Found & Müller, 1996; Müller et al., 1995; Liesefeld & Müller, 2019; Liesefeld et al., 2019, 2024; Sauter et al., 2018). Same-dimension distractors are therefore harder to suppress because they are defined in the task-critical dimension, whereas different-dimension distractors are less likely to compete for attentional selection.

This dimension-specific competition engages a distributed set of frontoparietal regions. Same-dimension distractors and changes within task-relevant visual dimensions recruit regions including the frontal eye fields (FEF), intraparietal and superior parietal regions (IPS/SPL), inferior frontal gyrus (IFG), and supplementary and pre-supplementary motor areas (SMA/pre-SMA; Pollmann et al., 2000; Serences et al., 2005; Corbetta et al., 2008; Müller et al., 2010). These regions have been implicated in attentional orienting and reorienting, spatial-priority updating, and control when stimulus-driven priority conflicts with current goals. Their response to same-dimension distractors therefore provides a neural index of the attentional demands generated when a salient distractor competes within the task-critical dimension. This makes the frontoparietal attention network a theoretically relevant locus for testing whether pre-stimulus state predicts subsequent distractor-evoked engagement.

Selection history adds a spatial layer to this architecture. When distractors repeatedly occur in one region, observers learn the regularity and show reduced costs for distractors at the frequent relative to rare locations (Goschy et al., 2014; van Moorselaar & Slagter, 2019; Wang & Theeuwes, 2018). The behavioral consequences of such location-specific learning may be especially evident when distractors generate substantial competition, as same-dimension distractors do. Different-dimension distractors can often be handled through dimension-specific priority settings, leaving less scope for an additional benefit from location-based probability cueing (Liesefeld & Müller, 2021; Sauter et al., 2018; Zhang et al., 2019). Learned spatial suppression is therefore not expressed independently of the current attentional context: its behavioral and neural consequences depend on the dimension-specific competition generated by the distractor (Allenmark et al., 2024).

A growing literature indicates that attentional processing depends on neural activity preceding stimulus onset. Electrophysiological studies have revealed temporally and spectrally specific forms of pre-stimulus modulation across cortical systems. Theta-rhythmic modulation of gamma activity in the visual cortex has been linked to the sequential sampling of attended locations (Landau et al., 2015). Alpha-band activity during the cue–target interval has also been shown to gate upcoming visual information, with sustained alpha desynchronization in dorsal intraparietal regions contralateral to the attended hemifield and alpha synchronization in ventrolateral extrastriate regions ipsilateral to the attended hemifield (Yang et al., 2024). More recently, beta- and gamma-band dynamics measured before target onset, during the cue–target interval, within attentional-network regions were shown to predict subsequent conscious reports of near-threshold targets (Spagna et al., 2026). Together, these studies indicate that neural states during the period preceding an upcoming target can shape subsequent attentional selection and perceptual outcomes, with evidence involving both sensory and frontoparietal attention systems.

Pre-stimulus fMRI findings provide complementary evidence at a slower temporal scale. In visual search, pre-trial activity in left middle frontal gyrus predicted momentary resistance to attentional capture (Leber, 2010), indicating that fluctuations in frontal control state can shape subsequent distractor processing. In sustained-attention tasks, attentional lapses were preceded by reduced activity in control-related regions and were followed by altered sensory and frontoparietal responses to the upcoming stimulus (Weissman et al., 2006). Together, these findings suggest that pre-stimulus state can shape both subsequent behavioral performance and the recruitment of frontoparietal attention systems by the upcoming event. This motivates testing whether a pre-stimulus state-sensitive fMRI measure predicts both behavioral responses and subsequent distractor-evoked frontoparietal activity, and whether these state–performance and state–network relationships vary across learned distractor-suppression contexts.

Two additional measures provide complementary views of the pre-stimulus state. The default-mode network (DMN) offers a network-level measure of intrinsic cortical activity: DMN fluctuations have been linked to internal mentation, task engagement, and vigilance (Raichle et al., 2001; Christoff et al., 2009; Konishi et al., 2015; Kamp et al., 2018), and weaker task-related DMN suppression has been associated with lapses and poorer performance (Raichle et al., 2001; Weissman et al., 2006; Konishi et al., 2015). Pupil size provides a peripheral psychophysiological comparison because fluctuations in pupil size index arousal- and vigilance-related variation and have been linked to locus-coeruleus-related BOLD activity and large-scale cortical dynamics (Murphy et al., 2014; Yellin et al., 2015).

The fMRI global signal (GS) offers a broad window onto this state. GS captures widespread trial-to-trial BOLD fluctuations and has been linked to vigilance, arousal-related dynamics, and autonomic physiological coupling (Wong et al., 2013; Chang et al., 2016; Liu et al., 2018; Bolt et al., 2025). Its interpretation is necessarily cautious. GS contains neural, vascular, motion-related, and other physiological contributions, and it does not isolate a single arousal mechanism. We therefore treat GS as a broad pre-stimulus cortical-state proxy and ask whether its trial-to-trial variation predicts subsequent performance and stimulus-evoked engagement of the frontoparietal attention network.

To test whether pre-stimulus state–performance mappings depend on learned distractor-suppression context, we reanalyzed an existing fMRI visual-search dataset (Shi et al., 2026). Participants searched for a uniquely tilted target bar while ignoring a singleton distractor defined either in the same dimension as the target, namely orientation, or in a different dimension, color. Spatial predictability was manipulated between groups by assigning the learned spatial-probability regularity to one distractor dimension. We refer to this as the biased dimension: distractors from this dimension appeared in a participant-specific frequent region on approximately 82% of the biased-dimension trials and in the opposite rare region on approximately 18%. Distractors from the other, unbiased dimension were distributed equally across the same two regions (50%/50%) and therefore carried no spatial-probability regularity. In the same-dimension spatial-bias (SS) group, the same-dimension distractor constituted the biased dimension, whereas in the different-dimension spatial-bias (DS) group, the different-dimension distractor constituted the biased dimension. The task therefore preserved the same stimulus structure across participants while placing the learned spatial regularity in different distractor-suppression contexts (**Figure 1**).

**Figure 1.**
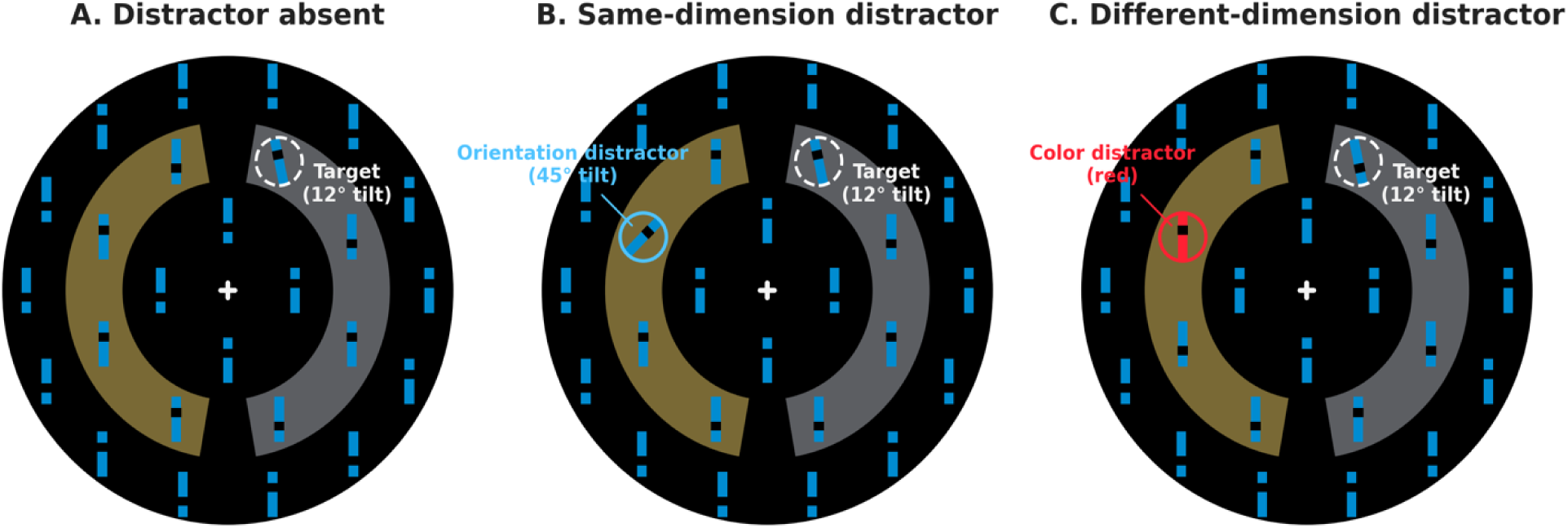
Task design schematic. Participants searched for a turquoise target bar tilted 12° from the vertical (in an array of upright turquoise non-target bars) and reported whether its notch pointed up or down; the position of the target and distractor (a distractor was present in 50% of trials) were restricted to the eight middle-ring locations. (**A**) Distractor-absent display. (**B**) Same-dimension distractor display with a turquoise orientation singleton tilted 45°; this constituted the biased dimension in the same-dimension spatial-bias (SS) group. (**C**) Different-dimension distractor display with an upright red color singleton; this constituted the biased dimension in the different-dimension spatial-bias (DS) group. Within the biased dimension, distractors appeared in a participant-specific frequent region in 82% of trials and in the opposite rare region in 18%; the unbiased dimension was distributed equally across the two regions (50%/50%). Yellow and gray sectors illustrate the frequent and rare regions, with left/right assignment counterbalanced across participants. Circles, labels, and shaded sectors are explanatory annotations and were not shown during the task.

This between-context design allowed us to test whether the behavioral consequences of pre-stimulus cortical state depend on the learned distractor-suppression context. Beyond the global signal, we focused the regional neural-state analyses on three systems with distinct theoretical roles. Attention and control networks were selected because frontal and parietal regions are consistently implicated in attentional orienting, priority updating, and resolving distractor competition (Corbetta et al., 2008; Müller et al., 2010). Early visual cortex was included because learned distractor-location regularities have been associated with reduced visual-cortical responses at frequent relative to rare distractor locations (Zhang et al., 2022). The default-mode network (DMN) was examined because its ongoing activity has been linked to internal mentation, task engagement, vigilance, and subsequent fluctuations in attentional performance (Raichle et al., 2001; Weissman et al., 2006; Christoff et al., 2009; Konishi et al., 2015; Kamp et al., 2018).

We therefore addressed four questions. First, do behavior and stimulus-evoked neural responses reproduce the expected dimension- and location-dependent pattern of distractor processing? Dimension-weighting accounts predict substantially stronger behavioral interference and frontoparietal engagement for same-dimension than different-dimension distractors, while prior fMRI work predicts reduced visual-cortical responses at frequent relative to rare distractor locations (Zhang et al., 2022). Second, how does pre-stimulus GS predict subsequent performance across the two learning contexts? And do regional neural activity and pupil size provide convergent evidence for the broader state–performance pattern? If GS reflects only a context-independent readiness state, its association with performance should be similar across groups; if instead its behavioral consequences depend on learned attentional context, the GS– performance mapping should differ between the same-dimension and different-dimension spatial-bias groups. Also, regional pre-stimulus BOLD activity and pupil size further clarify whether the state-behavioral effect is reflected across central and peripheral state-sensitive measures. Third, does pre-stimulus GS predict subsequent activity in independently defined attention/control, early visual, and default-mode regions, and does this modulation vary by distractor condition or learning context? This tests whether pre-stimulus state is expressed in subsequent regional cortical responsivity. Finally, we tested whether individual differences in GS-related neural attenuation are associated with the context-dependent GS–performance relationship, providing a link between pre-stimulus state, subsequent cortical responsivity, and behavior.

## Results

### 1 Functional Architecture of Distractor Interference and Spatial Learning

We first established the behavioral and stimulus-evoked neural architecture within which pre-stimulus state effects were evaluated. Group differences were tested through the relevant interaction, followed by group-specific simple effects.

### 1.1 Behavioral Distractor Interference and Spatial-Probability Learning

We used linear mixed-effects models to analyze task performance. Models entered phase (pre-scanner training vs. scanner), distractor dimension (same vs. different), distractor location (frequent vs. rare region), and group (SS vs. DS) as fixed effects, with subject as a random intercept.

Reaction times (RTs) declined continuously across pre-scanner training and scanning (**Figure 2A**). The joint model estimated a 13.3-ms decrease per block, *95% CI* [−14.8, −11.7], *p* < .001, with no reliable Group difference in the practice-related slope or scanner transition (both *p*s ≥ .299). Training and scanning were therefore treated as a continuous learning sequence.

**Figure 2.**
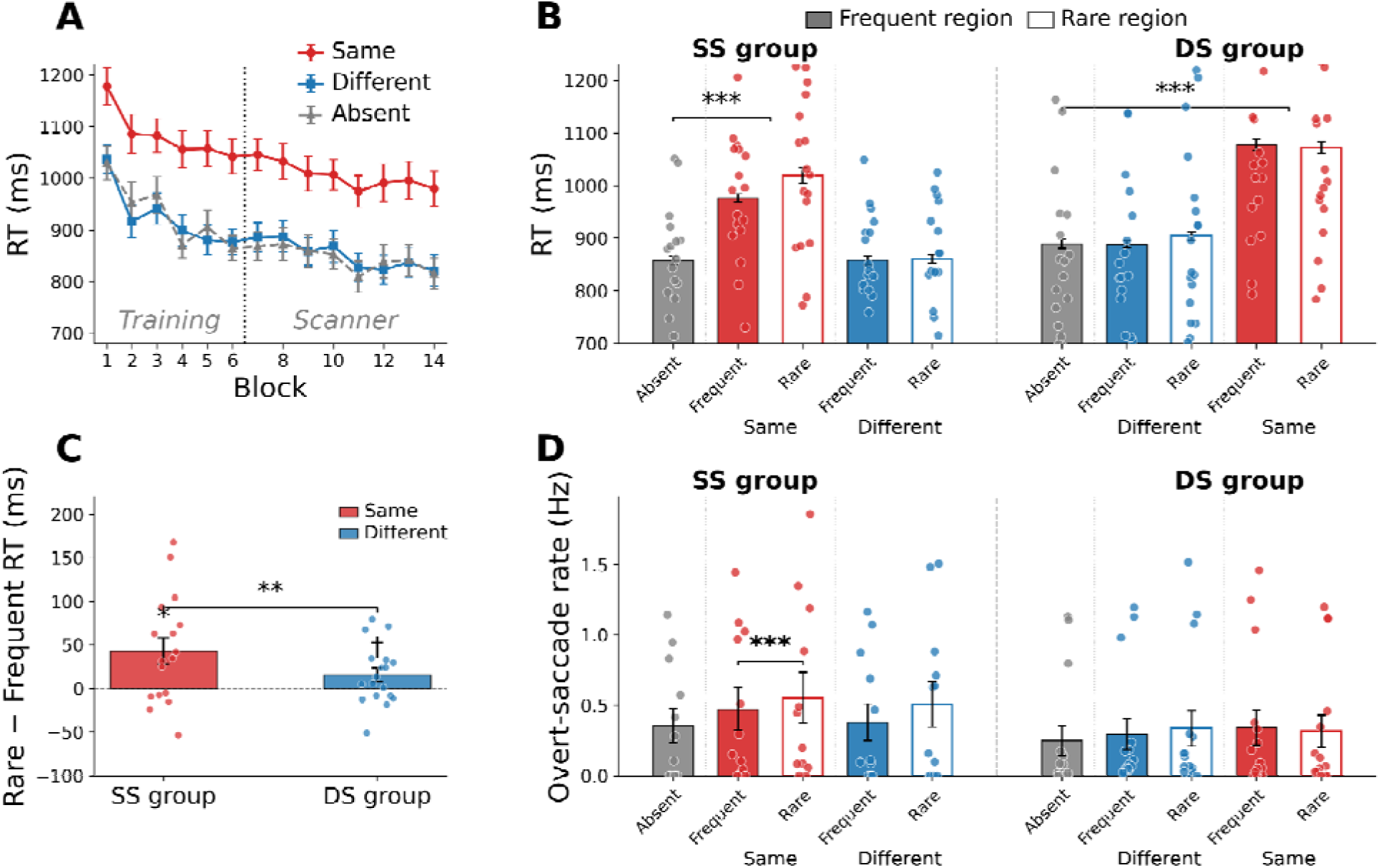
Behavioral results across the whole experiment. (**A**) Procedural RT improvement across training (blocks 1–6) and scanner (blocks 7–14) phases, plotted separately for same-dimension, different-dimension, and distractor-absent conditions. The phase boundary is marked by a dashed vertical line. (**B**) Mean RT by condition: Absent, biased-dimension distractors at Frequent and Rare locations, and unbiased-dimension distractors at the defined Frequent and Rare locations, shown separately for the SS (same-dimension spatial-bias) and the DS (different-dimension spatial-bias) group. (**C**) Location learning: Rare and Frequent RT difference for the biased dimension, by group. (**D**) Search-display overt-saccade rates as a function of distractor dimension and learned distractor region. Rates were calculated as the number of overt saccades divided by valid gaze time from display onset until the response or 1,000 ms, whichever occurred first. For all panels, individual subjects are shown as dots. Error bars = *SEM*. Significance markers denote Holm-adjusted (p)-values for the planned contrast family within each outcome; raw and Holm-adjusted (p)-values are reported in the Results. Significance markers: *** *p_Holm_* < .001, ** *p_Holm_* < .01, * *p_Holm_*< .05, †*p_Holm_* < .10.

The primary Group × Distractor-Dimension test was reliable (**Figure 2B**), β = +57.5 ms, 95% *CI* [38.1, 76.9], *p* < .001. There were substantial same-dimension RT costs in both groups: SS, β = +128.2 ms, *95% CI* [115.7, 140.8], DS, β = +185.7 ms, *95% CI* [170.9, 200.5], both *p_Holm_*s < .001. Different-dimension RT costs were negligible in both groups (SS: β = +1.2 ms, DS: β = +1.7 ms, both *p_Holm_*s > .99). The Group difference was not significant, β = +0.4 ms, *95% CI* [−18.8, 19.7], *p* = .964.

The Group × Location interaction, β = −29.3 ms, 95% *CI* [−50.9, −7.6], *p* = .008, indicated different spatial-bias patterns across groups. In the SS group, same-dimension distractors in the rare region slowed responses relative to same-dimension distractors in the frequent region (**Figure 2C**), Mean = 43.3 ms, 95% *CI* [11.7, 74.8], *t* (16) = 2.90, *p_Holm_* = .021. In the DS group, different-dimension distractors in the rare region showed no reliable slowing relative to different-dimension distractors in the frequent region, Mean = 15.5 ms, 95% *CI* [−1.3, 32.4], *t* (17) = 1.94, *p_Holm_*= .069. In contrast, for the unbiased dimension, the Group × Location interaction was not significant, β = −7.0 ms, 95% *CI* [−34.8, 20.8], *p* = .620. In both groups, distractors from the unbiased dimension showed no reliable rare-versus-frequent region RT difference (both *p_Holm_*s > .99; SS: *M* = 2.7 ms, 95% *CI* [−13.0, 18.5], DS: *M* = −5.2 ms, *95% CI* [−31.5, 21.1]).

Accuracy was high overall (90.6%). The Group × Distractor-Dimension interaction was significant, likelihood-ratio χ*²*(2) = 16.85, *p* < .001. Accuracy was lower for same-dimension than for distractor-absent trials in both groups (both *p_Holm_*s < .001). There was no difference in accuracy between different-dimension and distractor-absent trials in either group (both *p_Holm_*s > .93). The Group × Location interaction for accuracy was not significant (*p* = .128).

Together, these behavioral results established two features of the task architecture. First, same-dimension distractors produced substantial interference, whereas different-dimension distractors produced little reliable cost. Second, spatial-probability learning was expressed most clearly when the spatially biased distractor shared the target-defining dimension.

### 1.2 Pre- and Search-display Saccadic Measures

To determine oculomotor signatures of learned spatial prioritization under the central-fixation requirement, we examined eye movements before and after search-display onset. Specifically, we asked whether SS and DS differed in pre-display oculomotor preparation and whether learned Frequent versus Rare regions modulated search-display overt saccades. Microsaccades were defined as small eye movements of 0.10°–1.00°, whereas overt saccades were movements exceeding 2.00°.

During the final 500 ms before display onset, neither microsaccade nor overt-saccade rates differed reliably between SS and DS (all *p_Holm_*s ≥ .339). Thus, there was no evidence that the two groups entered the search display with different overt oculomotor preparation.

We next examined search-display eye movements. From display onset until the response or 1,000 ms, whichever occurred first, at least one overt saccade occurred on 24.3% and two or more saccades on 6.9% of valid trials. We treated overall overt-saccade rate as the primary search-display oculomotor measure, which showed a clear search-display modulation in SS (**Figure 2D**): On biased-dimension trials, the overt-saccade rate was higher for Rare than Frequent regions, rate ratio = 1.161, 95% *CI* [1.082, 1.247], *p_Holm_* < .001. The same numerical pattern was evident on unbiased-dimension trials, rate ratio = 1.152, 95% *CI* [1.001, 1.327], *p* = .049, though it did not survive correction, *p_Holm_*= .344. Averaged across dimensions, the SS Rare/Frequent rate ratio was 1.157, 95% *CI* [1.070, 1.250], *p_Holm_* = .002. Importantly, however, the location effect did not differ between biased and unbiased dimensions, *p*_Holm_ > .99, indicating that the overt-saccade modulation was not specific to the dimension carrying the learned regularity in the SS group. In contrast, the DS group showed no reliable overall Region effect, rate ratio = 1.005, 95% CI [0.972, 1.039], *p*_Holm_ > .99. The overall Region effect differed between groups: the Rare-versus-Frequent rate ratio was 15.1% larger in SS than in DS, 95% CI [5.8%, 25.2%], *p_Holm_* = .009.

First-saccade occurrence provided convergent but weaker evidence. On SS biased-dimension trials, a first overt saccade occurred less often for Frequent than Rare locations, *p_Holm_* = .023. However, the overall SS Rare − Frequent effect did not survive correction, *p*_Holm_ = .227. The corresponding DS effect was also not reliable, *p*_Holm_ = .571, nor was the SS–DS difference, *p*_Holm_ = .952. Distractor-absent trials further argued against a general group difference in oculomotor indices: SS and DS did not differ reliably in microsaccade rate, overt-saccade rate, or first-saccade occurrence (all *p*_Holm_*s* ≥ .723).

Finally, overt distractor capture was behaviorally consequential. Relative to trials without an eligible overt saccade, RTs on trials on which the first saccade landed on the distractor were 137.4 ms slower, β = 137.38 ms, 95% *CI* [83.48, 191.29], *p*_Holm_ < .001; this capture-related RT cost did not differ between groups, *p_Holm_* = .460.

Overall, the oculomotor results provided little evidence that learning was expressed as a pre-display directional bias in eye movements. Instead, they point to a search-display reduction in overt orienting to Frequent regions in SS. The nearly identical Rare/Frequent rate ratios in the biased and unbiased dimensions are also consistent with a learned spatial bias generalizing from the biased same dimension to the unbiased different dimension, in line with previous evidence for cross-dimensional transfer (Sauter et al., 2019; Liesefeld & Müller, 2020).

### 1.3 Stimulus-Evoked Neural Responses to Same- and Different-Dimension Distractors

Planned stimulus-evoked contrasts compared same-dimension and different-dimension distractor trials with distractor-absent trials and with each other. The whole-brain maps further localized the Distractor-Dimension effects across attention- and control-related regions (**Figure 3A**). For C1 (Same > Absent), significant clusters included the left frontal eye field (FEF) and bilateral intraparietal sulcus/superior parietal lobule (IPS/SPL), primarily within Dorsal Attention territory, together with the supplementary motor area/pre-supplementary motor area (SMA/pre-SMA), left inferior parietal lobule (IPL), and right inferior frontal gyrus (IFG), spanning Control and Salience/Ventral Attention territories. C2 (Different > Absent) produced no suprathreshold clusters. C3 (Same > Different) again showed significant effects in the SMA/pre-SMA area, left IPL, and right IFG. All reported clusters survived cluster-level FWE correction (**Supplementary Table S1**). No significant group differences emerged for any capture-related contrast (all Group × Contrast interactions *p* > .25). Additional exploratory whole-brain group comparisons are provided in supplementary materials (**Supplementary Table S3**).

**Figure 3.**
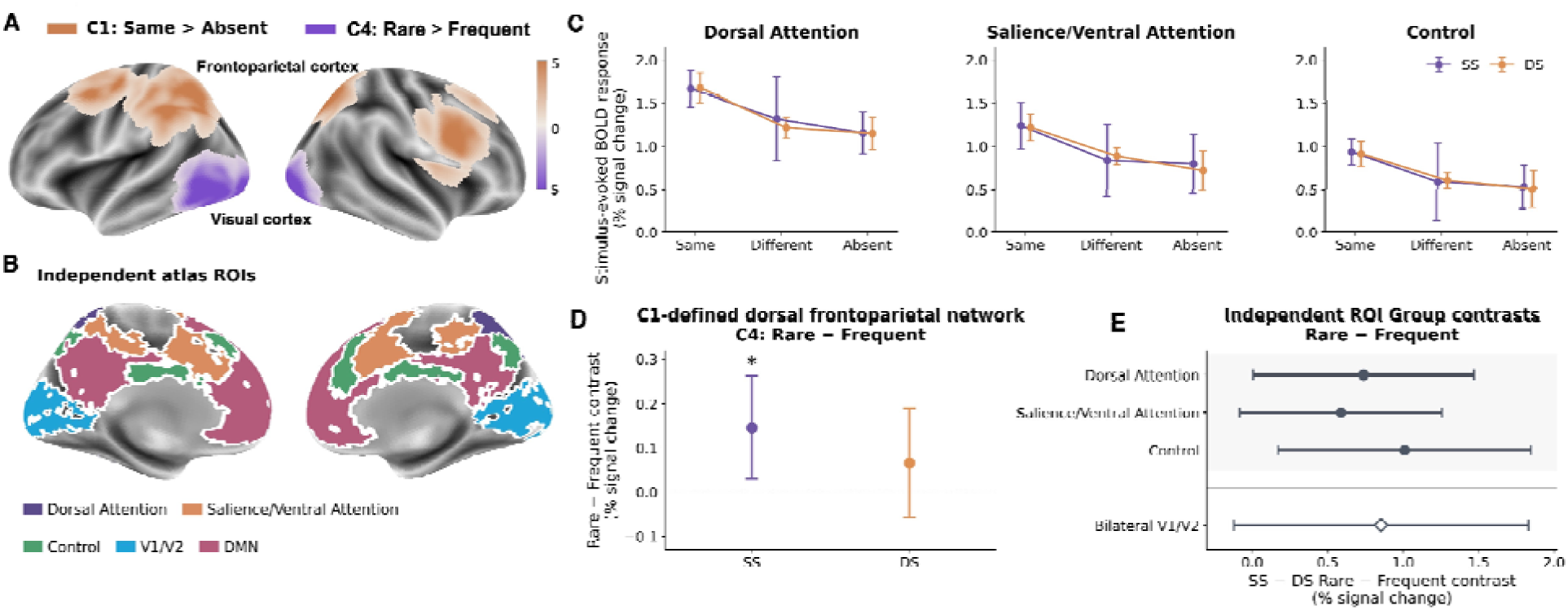
Stimulus-evoked neural responses and location-specific modulation. **(A)** Whole-brain activation maps for the Same > Absent contrast (C1; orange) and the Rare > Frequent contrast in the SS group (C4; purple), illustrating frontoparietal distractor-related activation and visual-cortical modulation by learned distractor location. **(B)** Independently defined regions of interest (ROIs) based on the Schaefer-400 parcellation with Yeo-7 network labels, including the Dorsal Attention, Salience/Ventral Attention, Control, and Default Mode networks, together with bilateral V1/V2 from the Juelich atlas. **(C)** Stimulus-evoked BOLD responses for same-dimension, different-dimension, and distractor-absent displays in the Dorsal Attention, Salience/Ventral Attention, and Control networks, shown separately for the same-dimension spatial-bias (SS) and different-dimension spatial-bias (DS) groups. **(D)** Rare-minus-Frequent contrast estimates (C4) in the C1-defined dorsal frontoparietal network for SS and DS. **(E)** Direct SS-minus-DS differences in the Rare-minus-Frequent contrast across independently defined Dorsal Attention, Salience/Ventral Attention, Control, and bilateral V1/V2 ROIs. Error bars indicate 95% confidence intervals.

This distribution broadly involved regions represented in the independently defined attention and control networks (**Figure 3B**). Based on the Schaefer-400 parcellation with Yeo-7 network labels, the Dorsal Attention Network comprises 46 parcels spanning posterior parietal cortex (PPC), frontal eye fields (FEF), and ventral precentral cortex; the Salience/Ventral Attention Network comprises 47 parcels spanning frontal operculum (FO) and insula, medialfrontal cortex (MFC), lateral prefrontal cortex (LPFC), parietal operculum, temporo-occipital/parietal cortex, and precentral cortex; and the Control Network comprises 52 parcels spanning parietal and temporal cortex, lateral and ventral prefrontal cortex (PFC), posterior-medial prefrontal cortex (pmPFC), precuneus, cingulate cortex, and left orbitofrontal cortex (OFC). Using these independently defined network masks as ROIs, same-dimension displays elicited higher activity than distractor-absent displays in Dorsal Attention (**Figure 3C**), β = 0.516%, 95% *CI* [0.256, 0.776], *t*(33) = 4.04, BH-FDR-*q* = .003, and Salience/Ventral Attention, β = 0.472%, 95% *CI* [0.124, 0.820], *t*(33) = 2.76, *q* = .042; the Control estimate did not survive correction, *q* = .086. Responses to different-dimension minus distractor-absent displays were not reliable in any network (all *q*s ≥ .600), and Same-minus-Different responses did not survive correction (all *q*s ≥ .101).

Consistent with our expectations and with the dimension-weighting account (e.g., Found & Müller, 1996; Liesefeld & Müller, 2019), the neural results converged with the behavioral findings: same-dimension distractors gave rise to stronger interference and recruited a distributed frontoparietal attention network, whereas different-dimension distractors produced substantially weaker behavioral and neural effects.

### 1.4 Location-Specific Neural Modulation

We next asked whether learned distractor location modulated stimulus-evoked activity within cortical systems involved in distractor processing. C4 was defined as Rare minus Frequent within each group’s probability-biased distractor dimension: Same-Rare minus Same-Frequent in SS and Different-Rare minus Different-Frequent in DS.

Within the prespecified dorsal frontoparietal network defined by the same-dimension capture contrast (C1; left FEF, left IPS/SPL, SMA/pre-SMA, and right SPL), C4 was positive in SS (**Figure 3D**), β = +0.146, 95% *CI* [+0.029, +0.262], *t* (16) = 2.66, *p* = .017, and smaller in DS, β = +0.066, 95% *CI* [−0.058, +0.190]*, t* (16) = 1.13, *p* = .276. However, the direct between-group comparison was not significant, SS − DS β = +0.080, 95% *CI* [−0.084, +0.243], *p* = .329. Direct SS − DS comparisons of the biased-dimension Rare − Frequent contrasts were positive in each independently defined attention/control network (**Figure 3E**): Dorsal Attention, β = +0.737%, 95% *CI* [+0.006, +1.469], *p* = .048, *q* = .086; Salience/Ventral Attention, β = +0.587%, 95% *CI* [−0.081, +1.256], *p* = .085, *q* = .086; and Control, β = +1.009%, 95% *CI* [+0.172, +1.846], *p* = .018, *q* = .073. The bilateral V1/V2 analysis also showed a difference but did not reach significance, SS − DS β = 0.854%, 95% *CI* [−0.121, 1.829], *q* = .086. Within SS, which showed the clearer behavioral spatial-learning effect, a secondary whole-brain analysis localized Rare > Frequent responses to bilateral occipital cortex and left lateral occipital complex (**Figure 3A**; **Supplementary Table S2**), consistent with previous fMRI findings on distractor-location learning (Zhang et al., 2022). Thus, location-related neural modulation was descriptively stronger in SS, particularly in frontoparietal and visual regions.

## 2 Pre-stimulus Cortical State Predicted Performance Differently Across Contexts

### 2.1 Pre-stimulus GS–RT Model

For each trial, we summarized GS as mean gray-matter BOLD amplitude during the 2 s before display onset, z-scored within run, and tested whether this pre-stimulus signal predicted RT after accounting for gradual practice or fatigue trends with log-transformed trial number.

Pre-stimulus GS predicted response speed differently across the two learned contexts (**Figure 4A**, left). In the distractor-present mixed-effects model, the GS × Group interaction was significant, β = −21.0 ms, 95% *CI* [−32.1, −10.0], *p* < .001, indicating that trial-by-trial GS fluctuations related to RT in opposite ways for the two groups. Distractor dimension did not reliably qualify this relationship: neither the GS × Distractor-Dimension interaction (β = +6.0 ms, *p* = .314, *BF* = 4.3) nor the GS × Distractor-Dimension × Group interaction (β = −3.7 ms, *p* = .687, *BF* = 5.6) reached significance. Higher GS predicted *slower* RT for same-dimension distractors in SS, β = +10.8 ms, *p_Holm_* = .006, and *faster* RT for different-dimension distractors in DS, β = −12.2 ms, *p_Holm_* = .002. The remaining two slopes did not reach the adjusted threshold (both *p_Holm_*≥ .177).

**Figure 4.**
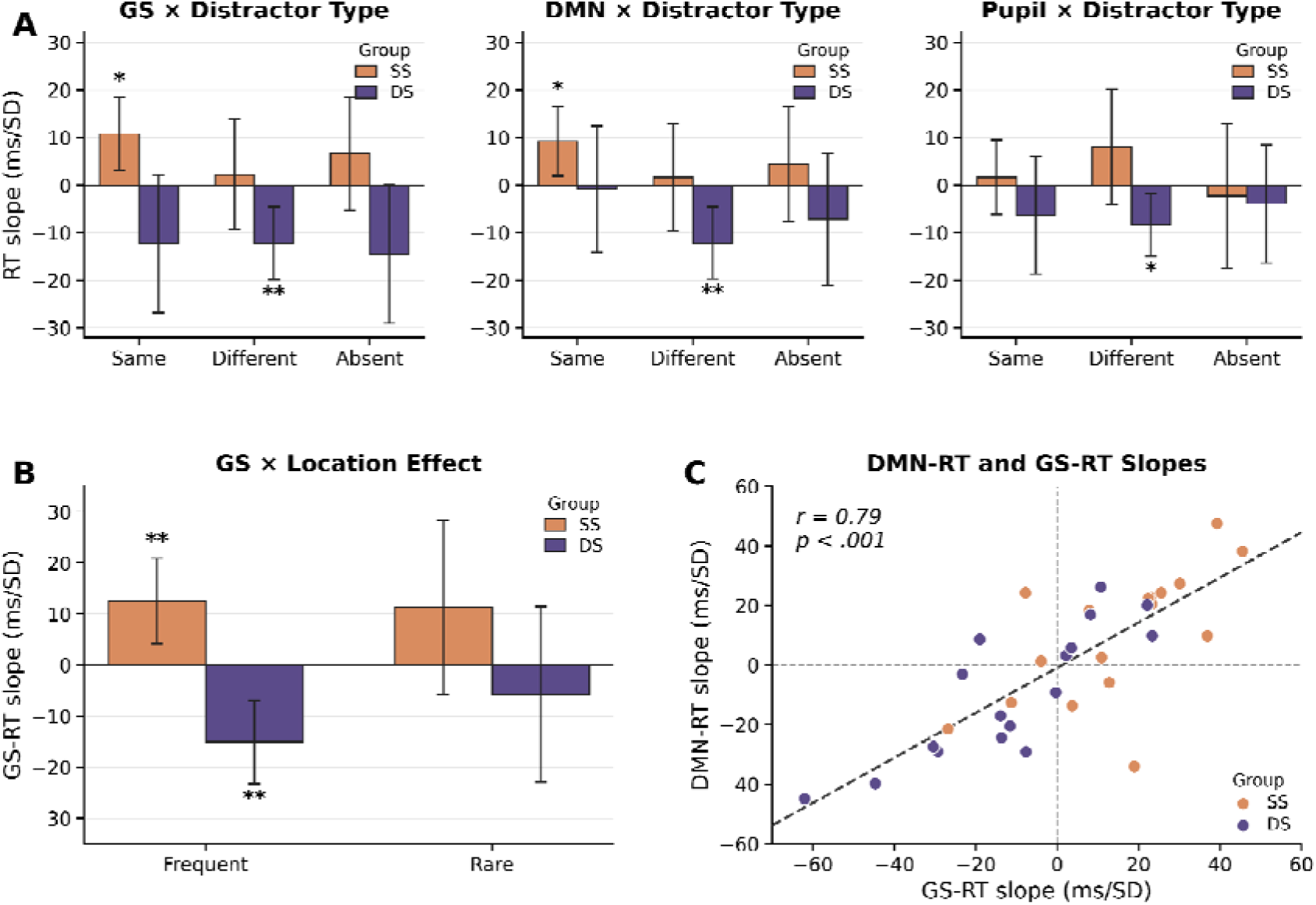
Pre-stimulus state-performance effects across GS, DMN, and pupil size. (**A**) Condition-specific RT slopes relating pre-stimulus global signal (GS; left), default mode network (DMN) activity (middle), and baseline pupil size (right) to response time in the different-dimension spatial-bias group (DS, blue) and same-dimension spatial-bias group (SS, red). (**B**) Location-specific GS-RT slopes for the biased dimension. (**C**) Subject-level DMN-RT slopes plotted against GS-RT slopes. Error bars in bar plots indicate *SEM*; * *p* < .05, ** *p* < .01.

The GS × Group × Location interaction was not significant, β = +10.6 ms, 95% *CI* [−16.1, 37.3], *p* = .438. Direct Rare-minus-Frequent slope contrasts were also not significant in SS, β = −1.2 ms, *p_Holm_* = .897, or DS, β = +9.3 ms, *p_Holm_* = .666. However, higher GS predicted *slower* RT in SS at the Frequent location, β = +12.5 ms, *p_Holm_* = .010, and *faster* RT in DS at the Frequent location, β = −15.1 ms, *p_Holm_*= .001 (**Figure 4B**); neither Rare-location slope was significant (both *p_Holm_* ≥ .393). Thus, the GS–RT relationship was context-dependent, differing between SS and DS, and was descriptively strongest at the frequent distractor location.

The GS-RT pattern remained reliable after controlling for previous trial type and for nuisance proxies related to motion, CSF signal, and white-matter signal; it also survived control for subject-level distractor-cost magnitude, arguing against a simple task-difficulty explanation. A quadratic model showed no reliable evidence for an inverted-U state-performance curve. Also, the oculomotor modulation explains only a small proportion of the context-dependent GS– performance relationship. Full robustness and quadratic follow-up statistics are reported in **Supplementary Results S4**.

### 2.2 Convergence Across Pre-stimulus ROI Activity and Pupil Measures

The identical pre-stimulus GS-RT model was applied separately to Dorsal Attention, Salience/Ventral Attention, Control, bilateral V1/V2, and DMN signals extracted from the same −2-to-0-s window on same-dimension, different-dimension, and distractor-absent trials. The DMN × Group interaction survived correction across this five-region family, β = −16.0 ms, 95% *CI* [−27.3, −4.7], *p* = .005, *q* = .027, whereas neither the DMN × Dimension interaction (β = +6.5 ms, *p* = .280) nor the DMN × Dimension × Group interaction (β = +3.8 ms, *p* = .679) reached significance. Condition-specific follow-ups reproduced the key GS pattern: in the DS group, higher DMN activity predicted *faster* responses for different-dimension distractors (β = −12.1 ms, *p* = .002), whereas in the SS group, higher DMN activity predicted *slower* responses for same-dimension distractors (β = +9.3 ms, *p* = .013). DMN and GS also converged at the level of individual differences. Participants with stronger GS-RT slopes tended to show stronger DMN-RT slopes in the same direction, *r* = .79, *p* < .001 (**Figure 4C**). The remaining Pre-stimulus regional activity × Group coefficients were not reliable: Dorsal Attention, β = −5.6 ms, 95% *CI* [−16.8, 5.6], *p* = .328, *q* = .491; Salience/Ventral Attention, β = −3.9 ms, 95% *CI* [−15.1, 7.3], *p* = .491, *q* = .491; Control, β = −4.1 ms, 95% *CI* [−15.3, 7.1], *p* = .472, *q* = .491; and V1/V2, β = −10.1 ms, 95% *CI* [−21.2, 1.0], *p* = .075, *q* = .188. The regional decomposition therefore revealed DMN only, but none of the other four atlas regions.

Nevertheless, pre-stimulus GS was positively correlated with all regional signals across distractor types (DAN *r* = .794–.813, Salience/VAN *r* = .754–.788, Control *r* = .760–.787, V1/V2 *r* = .728–.761, DMN *r* = .728–.763) and the strength of this coupling did not differ reliably between groups within any distractor type (all *q* ≥ .954).

Pupil size provided a more peripheral physiological comparison but yielded weaker evidence (**Figure 4A, right**). The SS-minus-DS pupil-RT slope contrast was positive but marginal, β = +8.64 ms/*SD*, 95% *CI* [-0.73, 18.02], *p* = .071, and participant-level pupil–RT slopes did not correlate positively with GS–RT slopes, *r* = −.124, *p* = .485.

Thus, the GS–RT relationship differed between the SS and DS groups and was mirrored by an analogous DMN–RT relationship, whereas the baseline pupil-size–RT relationship showed only marginal evidence of a corresponding group difference.

## 3 Pre-stimulus GS predicts subsequent cortical activity

### 3.1 Pre-stimulus GS modulation of attention/control-network activity

Across groups, higher pre-stimulus GS generally preceded weaker subsequent stimulus-evoked activity in three independently defined bilateral networks (**Figure 5A**). In the attention/control networks, all direct Group comparisons yielded *q*s ≥ .975. On same-dimension distractor trials, higher pre-stimulus GS was associated with weaker subsequent stimulus-evoked activity in the Dorsal Attention network, β = −0.410% signal change per SD increase in GS, 95% *CI* [−0.668, −0.152], *t* (33) = −3.24, *q* = .012, and the Control network, β = −0.441%, 95% *CI* [−0.676, −0.207], *t* (33) = −3.82, *q* = .005; the Salience/Ventral Attention network exhibited the same direction, β = −0.253%, 95% *CI* [−0.476, −0.029], *t* (33) = −2.30, *q* = .053. Different-dimension slopes were also negative across all three networks, β range = −0.437% to −0.345%, *q* range = .053–.080, as were absent-trial slopes, β range = −0.468% to −0.396%, all *q*s = .080, although none survived the nine-test correction.

**Figure 5.**
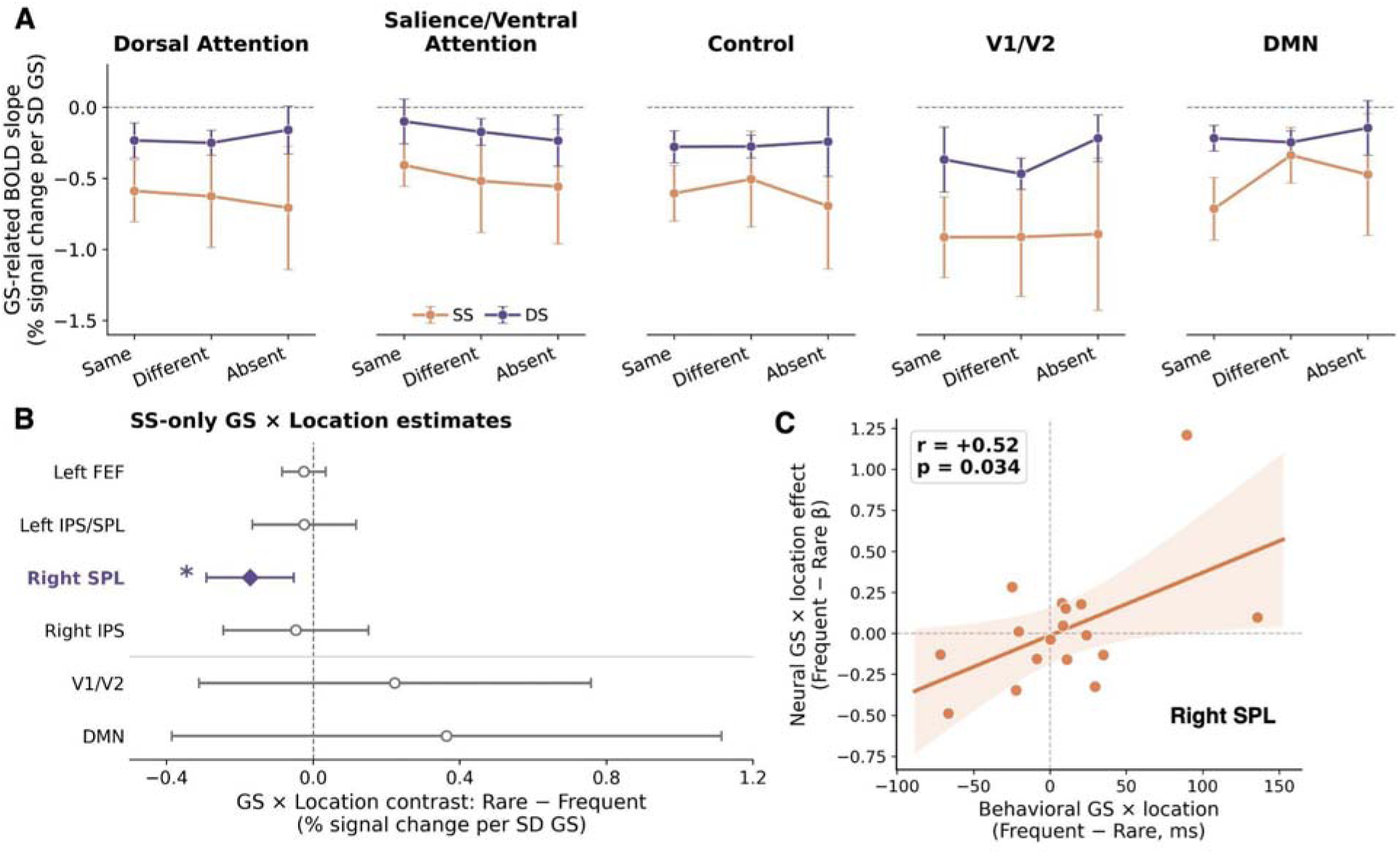
Pre-stimulus global-signal modulation across cortical systems and learned distractor locations. **(A)** Condition-specific relationships between pre-stimulus global signal (GS) and subsequent BOLD responses in independently defined Dorsal Attention, Salience/Ventral Attention, Control, bilateral V1/V2, and Default Mode Network (DMN) ROIs, shown separately for the same-dimension spatial-bias (SS) and different-dimension spatial-bias (DS) groups. Negative slopes indicate weaker subsequent stimulus-evoked activity at higher pre-stimulus GS. Error bars indicate 95% confidence intervals. **(B)** Exploratory GS × Location effects within the SS group across four Dorsal Attention regions—left frontal eye field (FEF), left intraparietal sulcus/superior parietal lobule (IPS/SPL), right superior parietal lobule (SPL), and right intraparietal sulcus (IPS)—together with bilateral V1/V2 and DMN. Estimates represent Rare-minus-Frequent differences in GS modulation; right SPL was the only ROI surviving BH-FDR correction. **(C)** Across SS participants, the behavioral GS × Location effect covaried with the corresponding neural GS × Location effect in right SPL, *r* = .52, *p* = .034. The line shows the fitted linear relationship with its 95% confidence interval. * indicates BH-FDR-corrected *q* < .05.

GS modulation extended to independently defined bilateral early visual cortex (V1/V2), defined as the union of cytoarchitectonic V1/BA17 and V2/BA18 masks from the Juelich maximum-probability atlas thresholded at 25%. Direct Group comparisons were also not reliable (all *q*s ≥ .583). Higher GS predicted weaker early visual activity on both same-, β = −0.641%, 95% *CI* [−1.018, −0.263], *t* (33) = −3.45, *q* = .006, and different-dimension trials, β = −0.690%, 95% *CI* [−1.130, −0.249], *t* (33) = −3.19, *q* = .006. The slopes did not differ reliably between the three types of trials (all *p*s > .70).

Condition-specific GS modulation also extended to the independently defined DMN ROI. Across participants, higher GS predicted weaker subsequent DMN activity on same-, β = −0.465%, 95% *CI* [−0.720, −0.211], *t* (33) = −3.72, *q* = .002, and different-dimension trials, β = −0.292%, 95% *CI* [−0.504, −0.079], *t* (33) = −2.79, *q* = .013, but not on distractor-absent trials, β = −0.309%, 95% *CI* [−0.783, 0.165], *t* (33) = −1.33, *p* = .194. The largest Group difference occurred on same-dimension distractor trials, for which the SS estimate was more negative than the DS estimate, difference = −0.496%, 95% CI [−0.981%, −0.011%], *t* (32) = −2.08, *p* = .045, *q* = .136. Different-dimension and distractor-absent trials showed no reliable Group differences (both *p*s ≥ .491).

### 3.2 Exploratory Location-Specific GS Modulation

Within the SS group, we tested whether GS modulation differed between rare and frequent locations on same-dimension distractor trials across four Dorsal Attention Network regions (left FEF, left IPS/SPL, right SPL, and right IPS) that showed the strongest attentional-interference responses (see Results Section 1.3). The composite GS × Distractor-Location effect was not significant, β = −0.068, 95% *CI* [−0.175, +0.039], *t* (16) = −1.34, *p* = .199. Right SPL (**Figure 5B**) was the only constituent region showing a reliable Rare-minus-Frequent difference after BH-FDR correction across the four ROIs, β = −0.172, *p* = .007, *q* = .030. Parallel analyses in independently defined V1/V2 and DMN ROIs showed no reliable GS × Distractor-Location effects, *ps* > .31. When correction was extended across all six ROIs, right SPL remained the only reliable effect (*q* = .045). GS modulation in right SPL was marginally positive at frequent locations, β = +0.130, 95% CI [0.00, 0.26], *t* (16) = 2.05, *p* = .057, *d* = 0.50, and negative though not significant at rare locations, β = −0.042, 95% *CI* [−0.20, 0.12], *t* (16) = −0.55, *p* = .591, *d* = −0.13. This neural pattern paralleled the behavioral GS × Distractor-Location follow-up. Individual differences in the behavioral and neural GS × Distractor-Location effects also covaried in right SPL (*r* = +.52, *p* = .034; **Figure 5C**). These findings provide tentative evidence that pre-stimulus state may influence the neural expression of learned spatial priority settings.

These results indicated that pre-stimulus cortical state predicted behavior in a context-dependent manner but showed broader neural effects. Higher GS generally preceded reduced activity across attention/control, visual, and DMN regions, with little evidence for group- or condition-specific neural modulation. Exploratory analyses suggested a possible location-specific effect in the right SPL.

## 4 ROI-Specific Coupling Between Neural Attenuation and Behavior

We next tested whether the between-context behavioral divergence is related to how strongly pre-stimulus GS attenuates subsequent regional activity. Stronger attenuation was defined as a more negative condition-specific GS–BOLD slope within each independently defined ROI, standardized separately by ROI and condition and sign-reversed. Separate cross-level models were fitted for Dorsal Attention, Salience/Ventral Attention, Control, V1/V2, and DMN within same-dimension, different-dimension, and distractor-absent trials.

Across distractor types and ROIs, the estimates showed a consistent directional pattern: stronger neural attenuation was associated with a more negative GS–RT relationship in DS and a more positive relationship in SS (**Figure 6**). However, for same-dimension trials, there was no evidence that attenuation moderated the GS–RT relationship differently between groups (all *q*s ≥ .407). Distractor-absent trials likewise showed no corrected Group differences (all *q*s ≥ .409).

**Figure 6.**
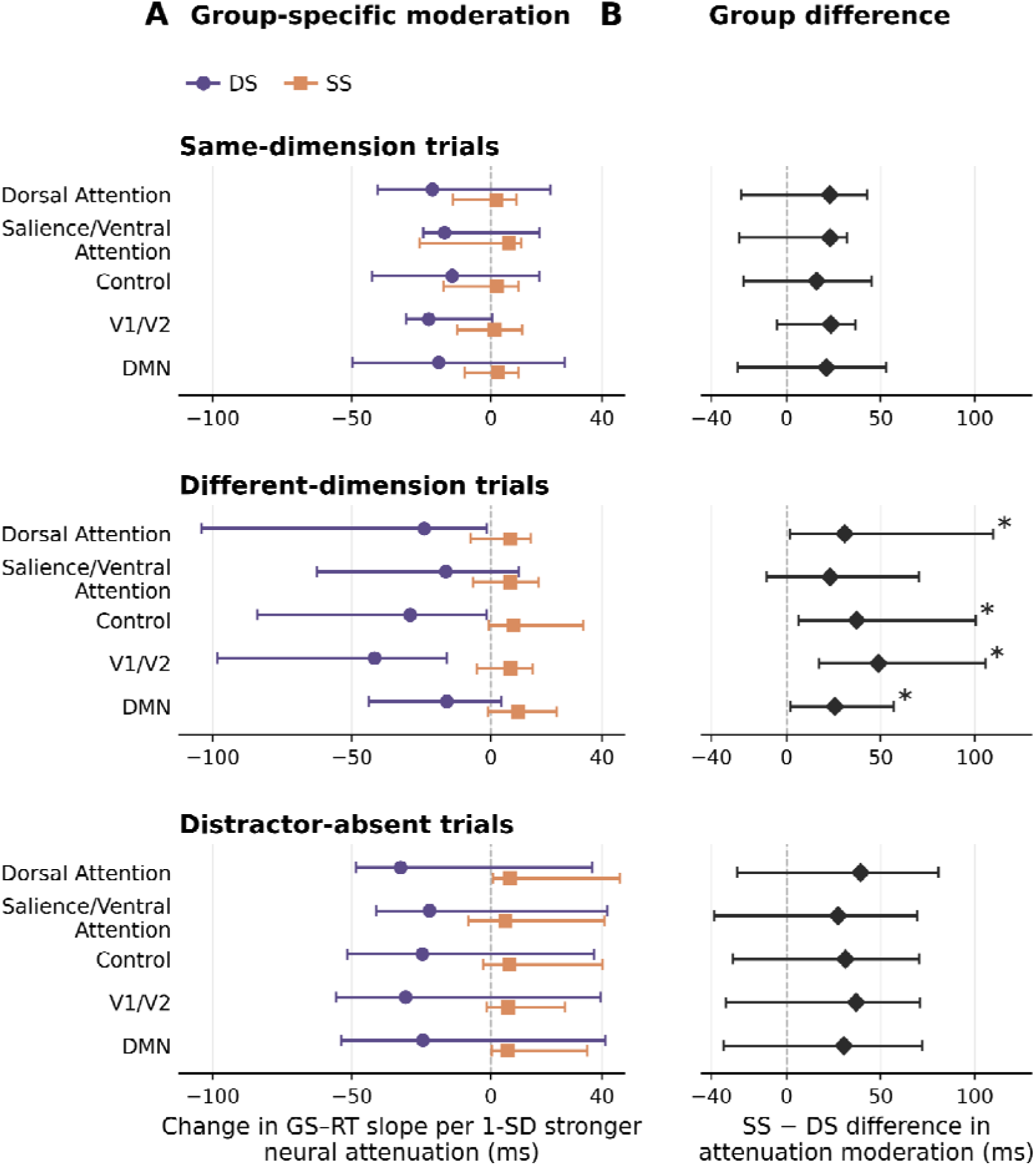
ROI-specific moderation of the relationship between pre-stimulus GS and RT by neural attenuation. (**A**) Within-group moderation estimates for DS and SS. Negative values indicate that participants with stronger GS-related neural attenuation showed a more negative trial-level GS–RT slope. (**B**) Direct SS-minus-DS differences in moderation. Points and horizontal lines show coefficient estimates and participant-bootstrap 95% confidence intervals from 5,000 resamples stratified by Group. \**q* < .05, survived BH-FDR correction across the five ROIs within that condition.

Different-dimension trials produced a broader positive SS-minus-DS moderation pattern. The direct Group differences were +30.9 ms in Dorsal Attention, 95% *CI* [1.7, 109.6], *p* = .032; +37.1 ms in Control, 95% *CI* [6.3, 100.5], *p* = .028; +48.8 ms in V1/V2, 95% *CI* [17.3, 105.6], *p*= .018; and +25.6 ms in DMN, 95% *CI* [2.1, 56.9], *p* = .020. All four survived BH-FDR correction across the five Different-trial ROIs, *q* = .039. The Salience/Ventral Attention estimate was +23.1 ms, 95% *CI* [−10.7, 70.2], *q* = .063. Across these ROIs, stronger attenuation was associated with a more negative GS–RT slope in DS, but little change in SS.

## Discussion

The present study asked whether the cortical state preceding a visual-search display shapes the subsequent expression of learned distractor suppression. Three findings define the overall pattern. First, the task reproduced the expected functional architecture of distractor interference: same-dimension distractors produced substantial behavioral costs and recruited distributed dorsal-attention, salience/ventral-attention, and control regions, whereas different-dimension distractors produced little behavioral or stimulus-evoked interference; spatial-probability learning was also behaviorally clearer when the biased distractor shared the target-defining dimension. Second, pre-stimulus global signal (GS) predicted subsequent performance in opposite directions across the two learned contexts, with higher GS preceding slower responses in same-dimension spatial-bias (SS) group but faster responses in different-dimension spatial-bias (DS) group. Third, this context dependence at the behavioral level contrasted with a broader neural effect: higher GS generally preceded weaker subsequent activity across attention/control networks, early visual cortex, and the DMN, though the neural GS estimates tended to be more negative in SS than DS across all distractor types. Finally, ROI-specific coupling analyses showed that stronger GS-related neural attenuation tended to accompany a more negative GS–RT relationship in DS and a more positive relationship in SS, with the strongest group difference on different-dimension trials. Together, these findings distinguish a context-dependent state–performance mapping from a broader modulation of subsequent cortical responsivity, while suggesting that individual differences in how pre-stimulus state propagates into sensory and attentional processing may contribute to the behavioral divergence between learning contexts.

The behavioral and stimulus-evoked results first establish the attentional architecture within which these state effects occurred. Same-dimension distractors strongly slowed search, consistent with the dimension-weighting account according to which the target-relevant orientation dimension receives greater attentional weight, making an orientation singleton particularly difficult to ignore (Found & Müller, 1996; Liesefeld & Müller, 2019; Liesefeld et al., 2019, 2024; Sauter et al., 2018). The corresponding fMRI responses were distributed across frontal and parietal territories associated with attentional orienting, priority updating, and control. Different-dimension distractors, by contrast, produced negligible behavioral interference and no reliable Different > Absent network response. This pattern is consistent with previous neural evidence that distracting information within a task-relevant dimension preferentially recruits frontal and parietal attention regions, including the frontal eye fields and intraparietal cortex, whereas information in a task-irrelevant dimension engages partly different and less strongly frontoparietal mechanisms (e.g., Wei et al., 2018). The convergence between behavioral and fMRI results therefore supports dimension-weighting notions: distractors that share the target-relevant dimension engage attentional-control systems more strongly than equally salient distractors defined in a task-irrelevant dimension.

Behaviorally, the Rare–Frequent difference was larger in SS than DS, indicating that location learning was expressed most clearly when the spatially predictive distractor also competed within the target-relevant dimension. Neural effects showed a similar pattern: Rare– Frequent contrast resulted in more positive activation in attentional networks in SS, and smaller in DS, though the direct between-group difference was weaker. Within SS, Rare > Frequent responses were localized at the whole-brain level to bilateral occipital and lateral occipital cortex – broadly consistent with previous evidence that statistical distractor learning attenuates processing at expected distractor locations (Wang & Theeuwes, 2018; Zhang et al., 2022; Goschy et al., 2014). These results therefore support location-dependent modulation within the SS group rather than the DS group.

Moreover, the saccadic results indicated a general location-dependent modulation of overt orienting in the SS group, with no comparable reliable modulation in DS. There was no reliable group difference in pre-display microsaccade or overt-saccade activity. After display onset, however, SS showed a lower overt-saccade rate when distractors appeared in the Frequent rather than Rare region, consistent across biased and unbiased dimensions. First-saccade occurrence showed the same general direction, although the effect was clearest for biased-dimension distractors. The similar Rare/Frequent rate ratios across dimensions suggest that the SS location effect was not confined to the dimension carrying the spatial regularity. This pattern broadly accords with evidence that statistical learning reduces oculomotor capture at frequent distractor locations (Sauter et al., 2021) and is consistent with previous evidence that spatial learning acquired with same-dimension distractors can transfer to distractors in another dimension (Sauter et al., 2019; Liesefeld & Müller, 2020).

Against this background, the central finding was that pre-stimulus GS predicted performance differently across learning contexts: higher GS was associated with slower responses in SS but faster responses in DS. The Group difference was strongest at the frequent location. This Group effect was not reliably modified by the distractor dimension, indicating a context-dependent state–performance relationship rather than an effect specific to the biased distractor dimension. Pre-stimulus DMN activity showed the same Group-dependent relationship with RT as GS, and participants’ DMN–RT and GS–RT slopes were strongly correlated.

Importantly, the same behavioral model applied to the other independently defined networks did not yield comparable corrected State × Group effects, identifying the DMN as the clearest regional correlate of the context-dependent behavioral pattern. This fits previous work linking pre-stimulus DMN state to fluctuations in task engagement and subsequent performance (Weissman et al., 2006; Kamp et al., 2018). Their convergence therefore suggests overlapping central BOLD-state variation.

This context dependence argues against treating GS as a simple scalar measure of favorable or unfavorable readiness. The GS–RT relationship differed in direction between SS and DS, but this group difference was not reliably qualified by the current distractor dimension or distractor location. If higher GS merely reflected better alertness, poorer alertness, or greater task difficulty, one would expect its relationship with RT to retain the same sign across groups, perhaps differing only in magnitude. Instead, the same broad state measure was associated with opposite behavioral consequences under the two learning regimes. One possibility is that the functional consequence of a global cortical state depends on the attentional configuration already established through experience. In SS, the spatial regularity concerned a distractor that generated strong competition within the target-relevant dimension and produced robust spatial learning.

The accompanying reduction in post-display overt orienting to the Frequent region was expressed across biased and unbiased dimensions, suggesting a relatively broad spatial-priority setting. In DS, the spatially predictable distractor was already handled efficiently through dimension-based selection, generated little interference, and produced no comparable reliable location-dependent modulation of overt orienting. These different learning contexts may therefore have established distinct but internally consistent attentional-control settings. Under this account, GS does not directly encode “suppression strength”; rather, its behavioral consequence depends on how the current cortical state interacts with the attentional settings required by the learned context. This interpretation is consistent with the broader pre-stimulus literature showing that ongoing cortical activity can alter subsequent attentional selection and perceptual outcome (Leber, 2010; Weissman et al., 2006), while extending that principle to learned distractor suppression. This possibility is also consistent with the distinction between proactive and reactive control, whereby proactive control involves maintaining task-relevant settings in advance, whereas reactive control is recruited more transiently in response to incoming interference (Braver, 2012).

In particular, Sauter and colleagues proposed that when distractor and target are defined in the same dimension, learned suppression operates at a relatively supra-dimensional spatial-priority level and can therefore influence processing of multiple signals within the frequent region, whereas different-dimension distractors can be handled more selectively through dimension-specific down-weighting (Sauter et al., 2018, 2019; Liesefeld & Müller, 2021). SS may have relied more strongly on a maintained spatial-priority setting whose consequences became expressed during search-display orienting, while DS could rely more efficiently on dimension-based filtering, reducing the need for location-specific spatial control. Thus, pre-stimulus GS may index a broad cortical state whose behavioral consequence depends on how that state interacts with the balance between maintained spatial-priority settings and stimulus-driven control required by the learned attentional context.

At the neural level, GS showed a broader and largely context-general pattern. Across independently defined Dorsal Attention, Salience/Ventral Attention, and Control networks, higher pre-stimulus GS generally preceded weaker subsequent BOLD responses, and the same negative relationship extended to early visual cortex and the DMN, though SS showed numerically more negative GS–BOLD estimates than DS across the ROIs. This pattern was broadly consistent across same-dimension, different-dimension, and distractor-absent displays.

Thus, the neural effect appears to reflect a broad modulation of subsequent cortical responsivity rather than a mechanism specific to the distractor dimension. The exploratory location-specific analysis nevertheless suggests that this broad state effect may interact locally with learned spatial priority. Within SS, the right superior parietal lobule (rSPL) was the only tested region showing a corrected GS × Location effect, and individual differences in this neural modulation covaried with the corresponding behavioral GS × Location effect. Given the role of rSPL in spatial priority and attentional orienting, this result provides a possible bridge between global pre-stimulus state and the spatial expression of learned attentional settings.

The ROI-specific coupling analysis directly links individual differences in GS-related neural attenuation to the context-dependent GS–RT relationship. Across distractor types, stronger GS-related neural attenuation tended to accompany a more negative GS–RT relationship in DS and a more positive relationship in SS, with the strongest pattern in Different-Dimension trials. This finding suggests that the behavioral consequences of pre-stimulus state may depend partly on how strongly that state propagates into subsequent sensory and attentional processing.

The present findings are also consistent with growing evidence that the fMRI global signal contains meaningful state-related information rather than reflecting nuisance variance alone. Global fluctuations have been linked to widespread neural activity, vigilance, arousal-related cortical and subcortical dynamics, basal-forebrain modulation, and autonomic coupling (Schölvinck et al., 2010; Wong et al., 2013; Chang et al., 2016; Liu et al., 2018; Gu et al., 2019; Turchi et al., 2018; Bolt et al., 2025). At the same time, GS also contains motion-, respiratory-, vascular-, and other non-neuronal contributions (Liu et al., 2017; Power et al., 2017, 2020). In this study, pupil size showed a weaker and only qualitatively similar group pattern, and pupil–RT slopes did not co-vary with GS–RT slopes across participants. Given the link between pupil fluctuations and arousal-related processes (Murphy et al., 2014; Yellin et al., 2015), this limited correspondence suggests that the GS effect cannot be reduced to a single arousal mechanism. We therefore interpret GS as a broad state-sensitive signal rather than a process-specific neural measure. The persistence of the GS–RT effect after controlling for previous trial type, motion, CSF, and white-matter signals indicates that the observed state–performance relationship was not readily explained by these nuisance proxies.

Several limitations nevertheless remain. The two learning contexts were manipulated between participants, with 17 participants per group, so the GS × Group effect cannot be interpreted as a within-person change in attentional-control state. In addition, the spatially biased distractor dimension was also presented more frequently than the unbiased dimension, meaning that group jointly varied the distractor dimension, spatial regularity, and relative exposure.

Although trial-history analyses suggested that the GS–RT pattern was not readily explained by recent or cumulative exposure alone, the present design cannot fully dissociate effects of learned spatial regularity from differential exposure. Finally, because respiratory and cardiac activity were not recorded directly, the physiological sources contributing to GS cannot be determined. Thus, the present results establish a robust relationship between pre-stimulus state, subsequent performance, and cortical activity, but do not isolate the specific learning, neural, or physiological processes generating that relationship.

In conclusion, pre-stimulus GS was associated with broadly reduced subsequent cortical responsivity, but its relationship with performance differed between the same-dimension spatial-bias (SS) and different-dimension spatial-bias (DS) contexts. Individual differences in GS-related neural attenuation also covaried differently with behavior across the two groups. These findings suggest that a broad pre-stimulus cortical state does not have a fixed behavioral consequence; rather, its effect on performance depends on the attentional-control context established by distractor dimension and spatial regularity.

## Methods

### Participants, Task, and Acquisition

We reanalyzed an existing fMRI visual-search dataset described in Shi et al. (2026). The public dataset included 35 healthy adults (21 female; mean age = 28.1 years, SD = 4.0, range = 22–41). One participant was excluded because of excessive head motion (>3 mm translation), yielding *N* = 34 for the present analyses (SS = 17, DS = 17). The groups did not differ in age, t(32) = −0.17, p = .869, or sex distribution, χ²(1) = 0.12, p = .724. Behavioral analyses included all 35 behaviorally valid participants, whereas fMRI analyses included 34 participants after excluding one participant for excessive head motion.

Participants searched for a uniquely tilted bar among upright bars and reported the target notch position. Each trial began with a jittered fixation interval (0.5 or 5.5 s, equal probability), followed by a 1000-ms search display; responses were accepted until 1900 ms after display onset, and the inter-trial interval was 3–4 s. Search displays contained 26 turquoise bars arranged in three rings around fixation (1.25°, 2.50°, and 3.75° eccentricity; viewing distance = 140 cm). The target tilted 12° left or right from vertical; this orientation difference to the upright nontarget items is sufficient to ensure efficient search, i.e.: target ‘pop-out’ (Liesefeld et al., 2016). Same-dimension distractors were orientation singletons: one nontarget tilted 45° from vertical – which, in terms of feature contrast with the upright non-targets, was perceptually more salient than the 12°-tilted target. Different-dimension distractors were color singletons: one upright nontarget appeared in red (RGB [255, 33, 51]). A separate pilot experiment matched the orientation and color (distractor) singletons for perceptual saliency using the procedure of Zehetleitner et al. (2012), ensuring that neither distractor was inherently more visually conspicuous. This did not imply equal attentional priority in the search task, because the orientation distractor shared the target-defining dimension whereas the color distractor was defined in a task-irrelevant dimension.

Trial types were intermixed randomly within blocks. Distractors were absent on approximately 17% of trials, belonged to the spatially biased distractor dimension on approximately 61%, and belonged to the spatially unbiased distractor dimension on approximately 22%. Within the biased dimension, distractors appeared in a participant-specific frequent lateral region on 82% of biased-dimension trials and in the opposite rare lateral region on 18%; distractors from the unbiased dimension were distributed equally across the same two regions (50%/50%). In the same-dimension spatial-bias group (SS; N = 17), same-dimension distractors constituted the biased dimension, whereas in the different-dimension spatial-bias group (DS; N = 17), different-dimension distractors constituted the biased dimension. Assignment of the frequent region to the left or right visual field was counterbalanced across participants (**Figure 1**). Participants completed six pre-scanner training blocks (276 trials) followed by two scanner runs of 184 trials each.

Participants were instructed to maintain central fixation by looking at the middle of the display, but fixation was not enforced online and the display was not gaze-contingent. Eye position was recorded continuously with an EyeLink 1000 system (SR Research; 1 kHz sampling) and evaluated post hoc. Among imaging participants with valid fixation-period gaze, 91.5% within 3° of the display center. During the full 1-s search display, a saccade larger than 2° occurred on 24.2% of trials, whereas only 5.4% of trials contained such a saccade within the first 300 ms. These data indicate that gaze was largely central during fixation and early search, while overt eye movements occurred on a subset of trials later in the display. Stimulus presentation and response recording used Matlab with Psychophysics Toolbox (Brainard, 1997; Pelli, 1997).

Functional data were acquired on a 3T Siemens Prisma scanner with a 64-channel head coil using multiband EPI (TR = 1000 ms, TE = 30 ms, flip angle = 60°, multiband factor = 4, 60 slices, 2.5-mm isotropic voxels, FOV = 210 mm). A high-resolution T1-weighted MPRAGE image (TR = 2300 ms, TE = 2.32 ms, TI = 900 ms, 1-mm isotropic voxels) was acquired for anatomical registration.

### Preprocessing

Preprocessing used fMRIPrep 23.1.0 (Esteban et al., 2019), including motion correction, slice-timing correction, susceptibility-distortion correction, T1 registration, and normalization to MNI152NLin2009cAsym space. Functional images were resampled to 3-mm isotropic voxels. Unsmoothed fMRIPrep-preprocessed BOLD images were used for the regional pre-stimulus signals; response GLMs were smoothed at 6-mm FWHM. First-level models included six motion parameters, their temporal derivatives, and mean CSF and white-matter (WM) signals as confound regressors. Volumes with framewise displacement > 0.5 mm were flagged for quality control but retained to preserve temporal continuity.

### Region of Interest (ROI)

The independent network ROIs were based on the Schaefer et al. (2018) 400-parcel cortical parcellation at 2-mm source resolution with Yeo et al. (2011) seven-network labels. All bilateral parcels carrying the corresponding label were combined within each mask: Dorsal Attention comprised 46 parcels, Salience/Ventral Attention 47 parcels, Control 52 parcels, and DMN/Default 91 parcels. The atlas was resampled by nearest-neighbor interpolation to the 65 × 77 × 60 first-level analysis grid (3 × 3 × 3.3 mm). The resulting fixed masks contained 3,784 Dorsal Attention, 3,857 Salience/Ventral Attention, 5,016 Control, and 8,591 DMN voxels.

The independent bilateral early-visual ROI was the union of the GM Visual cortex V1 BA17 and GM Visual cortex V2 BA18 labels from the Juelich maximum-probability atlas thresholded at 25% (maxprob-thr25-1mm; Eickhoff et al., 2005). The V1 and V2 masks were resampled to the first-level grid using nearest-neighbor interpolation and combined into a 2,119-voxel V1/V2 mask. Thus, the five independent ROIs used for the regional state decomposition and subsequent neural-response analyses were Dorsal Attention, Salience/Ventral Attention, Control, bilateral V1/V2, and DMN (**Figure 3B**). These ROI masks were used across the stimulus-response, pre-stimulus regional-state, GS-modulation, and neural attenuation–behavior analyses.

A separate exploratory localization family used five 8-mm-radius spherical ROIs centered on peaks from the C1 Same-minus-Absent response map (**Figure 3A**): left FEF (MNI −37, −1, 60), left IPS/SPL (−34, −58, 50), SMA/pre-SMA (6, 21, 44), right SPL (18, −67, 53), and right IPS (33, −52, 50). These ROIs were selected from stimulus-evoked activity and were therefore not independent tests of C1. The centers of left FEF, left IPS/SPL, right SPL, and right IPS fell within the Schaefer/Yeo Dorsal Attention network (DAN) and were averaged to form the four-region DAN-centered capture composite.

### Pre-stimulus State Measures

Pre-stimulus global signal (GS) was computed as the mean BOLD signal across a gray-matter mask restricted to cortical and subcortical gray matter. For each trial, we averaged GS over the 2-s window immediately preceding display onset (2 volumes at TR = 1 s) and z-scored values within run. Because GS can contain neural and physiological variance, we tested whether trial-level GS covaried with framewise displacement, CSF signal, and WM signal (see S**upplementary Results S4**). No direct respiratory or cardiac recording was available, so residual physiological contributions could not be modeled explicitly (Power et al., 2020; Bolt et al., 2025).

Pre-stimulus regional signals were extracted using a common pipeline across all ROIs: Dorsal Attention, Salience/Ventral Attention, Control, and Default Mode networks, as well as bilateral V1/V2. For each ROI, the regional time series was linearly detrended and band-pass filtered at 0.01 - 0.10 Hz within run, standardized within run, and averaged over the two volumes immediately preceding display onset. The resulting trial-level signals were entered separately into the same behavioral model.

Pupil data were extracted from the scanner-phase EyeLink files and preprocessed using a pipeline adapted from de Gee et al. (2021) and Knapen et al. (2016). Samples with implausibly small pupil values (<100 arbitrary units) were treated as invalid. Blink periods were identified from the EyeLink event annotations, padded by 150 ms before blink onset and after blink offset, and linearly interpolated. The interpolated pupil trace was then low-pass filtered with a third-order Butterworth filter at 10 Hz. To reduce residual eye-movement-related pupil artifacts, blink and saccade onset regressors were convolved with a canonical pupil impulse-response function and regressed from the filtered trace, together with their temporal derivatives. For each trial, pre-stimulus pupil size was defined as the mean cleaned pupil signal during the 500-ms baseline window immediately preceding trial onset. This trial-level baseline measure was then standardized within participants, yielding a z-scored pre-stimulus pupil-size variable for subsequent analyses.

Pre-stimulus BOLD-based measures (GS and regional signals) were averaged over the 2 s immediately preceding display onset. Given the slower temporal dynamics and lower sampling rate of the BOLD signal, this window provided a relatively stable estimate of the local pre-stimulus cortical state while remaining close to stimulus onset. In contrast, pupil diameter was sampled at much higher temporal resolution and exhibits faster fluctuations; pre-stimulus pupil size was therefore defined over the final 500 ms before display onset to capture the immediate tonic pupil state while limiting averaging over earlier within-trial variation (Knapen et al., 2016; de Gee et al., 2021). These measures were not intended to provide temporally equivalent estimates of the same underlying physiological process.

### Behavioral and State-Performance Models

#### Practice and distractor-interference

Behavioral RT analyses used all 35 behavior-valid participants (17 SS, 18 DS), retained correct trials with RTs between 200 and 3000 ms; one-sided high-RT outliers more than 3 SD above each subject × dimension mean were excluded. The linear mixed-effects models used participant-specific random intercepts: RT ∼ block + block × Group + scanner phase + scanner phase × Group + Same + Same × Group + Different + Different × Group. Blocks were numbered continuously from 1 to 14, scanner phase was coded 0 for training and 1 for scanning, Same and Different were indicators relative to Absent trials, and Group was coded 0 for SS and 1 for DS. This model provided the linear practice slope, scanner-transition test, and same- and different-dimension interference cost reported in Results **1.1**. To assess whether overt eye movements influenced the behavioral effects, we repeated the RT analyses after excluding trials containing a >2° saccade (**Supplementary Results S5**).

#### Spatial-probability learning

Spatial-probability learning was tested across training and scanning in each Group’s biased distractor dimension: Same-dimension distractor trials in SS and different-dimension distractor trials in DS. A participant-random-intercept mixed model fitted RT ∼ block + Location × Group, with Frequent as the Location reference and SS as the Group reference. The Location × Group coefficient directly compared the Rare-minus-Frequent effect between SS and DS. Within-group Rare-minus-Frequent effects were then computed from participant-level mean differences and Holm-corrected across the two groups.

#### Accuracy

Scanner-phase accuracy was analyzed with participant-random-intercept binomial generalized linear mixed-effects models. The condition model fitted Accuracy ∼ Distractor Dimension × Group; the location model fitted Accuracy ∼ Location × Group within each Group’s biased distractor dimension. Likelihood-ratio tests compared each interaction model with its additive counterpart. Group interactions were tested before group-specific simple effects.

#### Oculomotor measures

We excluded EyeLink-detected saccades if they overlapped a blink, occurred within 50 ms of a blink, or had start or end coordinates outside the screen. The remaining events were classified as microsaccades when their amplitudes were 0.10°–1.00° and their durations were 6–100 ms. Events with amplitudes greater than 2.00° and durations of at least 6 ms were classified as overt saccades. Events with amplitudes greater than 1.00° but no greater than 2.00° were not included in either category. Pre-display activity was quantified during the final 500 ms before display onset on long-fixation trials with at least 50% valid gaze; event rates were expressed per second of valid gaze time. Search-display overt-saccade rates were calculated from display onset to the earlier of the manual response or 1,000 ms. Because event counts are discrete and the amount of usable gaze time varies across trials, we analyzed trial-level counts with Poisson generalized estimating equations. The GEEs clustered repeated trials within participants and used an exchangeable working correlation. We entered the log of trial-specific valid-gaze duration as an offset, which adjusted each count for its observation time and allowed the model to estimate event rates per second of usable gaze. Fixed effects comprised Group × Distractor Dimension × Region and centered log-transformed trial number.

Complementary first-saccade analyses tested whether an overt saccade began 80–800 ms after display onset and before the response. We calculated participant-level Rare-minus-Frequent occurrence contrasts and tested the corresponding Dimension and Group interactions. Endpoint analyses further required the first saccade to begin within 1.5° of fixation and end within 1° of the target or distractor. Distractor-absent trials were analyzed separately for early microsaccade rate (0–300 ms), overt-saccade rate, and first-saccade occurrence.

#### GS-RT model

The trial-level state–performance analyses used the canonical *N* = 34 sample. Pupil trials were merged with the scanner behavioral data by subject, session, and trial number. We applied an additional pupil-specific outlier filter within each subject × dimension cell, excluding trials whose standardized pupil size deviated by more than 3 SD from that cell’s mean. These procedures resulted in 5,164 same-dimension, 5,163 different-dimension, and 2,173 distractor-absent trials.

The primary GS model tested distractor-present RT as a function of log-transformed trial number, pre-stimulus GS, distractor dimension, and group: RT ∼ log(1 + trial number) + GS × Dimension × Group + (1|Subject). Log-transformed trial number controlled for practice, fatigue, and gradual learning across the scanner session. The critical terms were GS × Group, GS × Dimension, and GS × Dimension × Group. The location follow-up model was RT ∼ log(1 + trial number) + GS × Group × Location + (1|Subject). Condition-specific GS–RT slopes were estimated as simple effects from the joint Group model and Holm-corrected across the prespecified slope family. Parallel models replaced GS with pre-stimulus ROI regional activity or baseline pupil size. A trial-level Pearson correlation between GS and regional activity was computed separately for each participant across both runs. Correlations were Fisher-z transformed for group-level inference and comparison between SS and DS. And its mean and 95% confidence interval were back-transformed to the r scale. Robustness models added previous trial type, subject-level distractor-cost magnitude, framewise displacement, CSF signal, and white-matter signal (**Supplementary Results S4**). A quadratic model tested whether GS followed a non-linear state-performance function by adding GS² and its interaction with the group (S**upplementary Results S4**).

### Stimulus-Evoked BOLD Responses and Pre-stimulus GS Modulation

First-level GLMs were estimated with nilearn 0.10.1 (Abraham et al., 2014). Event regressors modeled same-dimension, different-dimension, and distractor-absent trials and were convolved with the canonical hemodynamic response function, accompanied by condition-specific pre-stimulus GS modulators. Stimulus-evoked contrasts established the functional architecture of distractor processing: C1 compared same-dimension distractor responses with distractor-absent trials, C2 compared different-dimension distractor responses with distractor-absent trials, and C3 directly compared same-with different-dimension distractors. The location-learning contrast was C4 (Rare > Frequent) within the spatially biased distractor dimension.

Primary ROI inference used the independently defined Dorsal Attention, Salience/Ventral Attention, and Control ROIs described above. Whole-brain analyses used a voxelwise threshold of *z* > 3.09 (*p* < .001 uncorrected) with cluster-level FWE correction (*p* < .05) in the second-level GLM. Group × Contrast interactions tested whether effects differed by learning context using run-averaged contrast images. To account for condition differences in saccade rate, a sensitivity analysis modeled each observed >2° display-period saccade at its onset as an HRF-convolved nuisance event, retaining the original regressors in participants with sufficient scanner gaze coverage (**Supplementary Results S5**).

Condition-specific GS slopes quantified the change in stimulus-evoked BOLD response (percentage signal change) associated with a 1-SD increase in within-run standardized pre-stimulus GS. Negative slopes indicated that higher pre-stimulus GS predicted a weaker subsequent response—that is, GS-related neural attenuation—whereas positive slopes indicated enhancement. For each participant, voxelwise GS-modulator coefficients were averaged within each atlas-defined ROI separately for each run and condition, and the resulting ROI coefficients were then averaged across the two runs. Direct contrasts tested whether GS-related modulation differed between same- and different-dimension, same-dimension and distractor-absent, and different-dimension and distractor-absent conditions.

### Neural Attenuation–Behavior Coupling Models

To test whether the context-dependent GS–performance relationship was related to individual differences in how pre-stimulus GS modulated subsequent regional activity, we examined the coupling between GS-related neural attenuation and trial-level GS–RT effects. For each ROI and condition, we first quantified each participant’s GS-related neural attenuation using the participant-specific GS–BOLD slope. These coefficients were standardized across participants and sign-reversed so that higher values indicated stronger attenuation of subsequent BOLD activity following higher pre-stimulus GS. We then tested whether this neural attenuation was related to the trial-level GS–RT relationship. Separate models were fitted for the Dorsal Attention, Salience/Ventral Attention, Control, V1/V2, and DMN ROIs within same-dimension, different-dimension, and distractor-absent conditions. Each model predicted RT from pre-stimulus GS, Group, neural attenuation, and their interactions, while controlling for trial number and run. DS served as the reference group. The critical GS × attenuation × Group interaction tested whether the relationship between neural attenuation and the GS–RT slope differed between DS and SS. These analyses treated the participant-level neural attenuation estimates as observed moderators. We additionally used participant-level bootstrap procedures to estimate confidence intervals for the moderation effects and to test the Group difference. Participant-level pairs bootstrapping was used to obtain confidence intervals for the moderation effects.

Participants were resampled with replacement within each Group while retaining all trials from each selected participant, and 5,000 resamples were used to derive percentile confidence intervals for the DS and SS estimates and their difference. Significance of the GS × attenuation × Group interaction was assessed with a restricted wild-cluster bootstrap-t, using 5,000 resamples. Benjamini–Hochberg correction was then applied to the resulting Group-difference p values across the five ROIs within each condition and across all 15 ROI × condition tests.

### Multiple-Comparison Correction

Multiple-comparison correction was defined according to the relevant inferential family. Holm correction was applied to small sets of planned follow-up contrasts, including behavioral, oculomotor, and condition-specific GS–RT comparisons. Benjamini–Hochberg false discovery rate (BH-FDR) correction was used for families of correlated ROI-level tests, whereas whole-brain analyses used cluster-level family-wise error (FWE) correction.

## Acknowledgments

This work was supported by German Research Foundation DFG grants CH3093/1-1 and SH 166/10-1, awarded to SC and ZS, respectively, and was carried out using the NICUM Siemens Prisma scanner supported by DFG grant INST 86/1739-1 FUGG.

## Scientific Transparency Statement

### DATA

All raw and processed data supporting this research are publicly available. The source fMRI visual-search dataset analyzed in the present study is publicly available at https://doi.org/10.12751/g-node.zfrem9.

### CODE

All analysis code supporting this research is publicly available at https://osf.io/sg4fw/.

### MATERIALS

This research did not make use of any materials to generate or acquire data. The present study is a reanalysis of previously collected data.

### DESIGN

This article reports how the authors determined the sample size, all data exclusions, all data inclusion and exclusion criteria, and whether inclusion and exclusion criteria were established prior to data analysis.

### PRE-REGISTRATION

No part of the study procedures was pre-registered in a time-stamped, institutional registry prior to the research being conducted. No part of the study analysis plans was pre-registered in a time-stamped, institutional registry prior to the research being conducted.

For full details, see the *Scientific Transparency Report* in the supplementary data to the online version of this article.

## Funding

This work was supported by the German Research Foundation (Deutsche Forschungsgemeinschaft, DFG) through grants **CH 3093/1-1** and **SH 166/10-1**. Data acquisition was carried out using the NICUM Siemens Prisma scanner, supported by DFG grant **INST 86/1739-1 FUGG**.

## Competing interests

The authors declare no competing interests.

## Declaration of Generative AI and AI-Assisted Technologies in the Manuscript Preparation Process

During the preparation of this work, the authors used Claude and Codex AI tools to polish the language and to tidy and optimize Python analysis code. After using these tools/services, the authors reviewed and edited the content as needed and took full responsibility for the content of the published article.

## Notes

### Competing Interest Statement

The authors have declared no competing interest.

### Summary of Updates

Independent ROI analyses have been added to support the results.

## References

Abraham, A., Pedregosa, F., Eickenberg, M., Gervais, P., Mueller, A., Kossaifi, J., Gramfort, A., Thirion, B., & Varoquaux, G. (2014). Machine learning for neuroimaging with scikit-learn. Frontiers in Neuroinformatics, 8, 14. 10.3389/fninf.2014.00014

Allenmark, F., Stanković, M., Müller, H. J., & Shi, Z. (2024). Dynamic suppression of likely distractor locations: Task-critical modulation. Visual Cognition, 32, 1045–1066. 10.1080/13506285.2024.2393467

Bolt, T., Wang, S., Nomi, J. S., Setton, R., Gold, B. P., Frederick, B. deB, Yeo, B. T. T., Chen, J. J., Picchioni, D., Duyn, J. H., Spreng, R. N., Keilholz, S. D., Uddin, L. Q., & Chang, C. (2025). Autonomic physiological coupling of the global fMRI signal. Nature Neuroscience, 28, 1327–1335. 10.1038/s41593-025-01945-y

Brainard, D. H. (1997). The Psychophysics Toolbox. Spatial Vision, 10, 433–436. 10.1163/156856897X00357

Braver, T. S. (2012). The variable nature of cognitive control: A dual mechanisms framework. Trends in Cognitive Sciences, 16, 106–113. 10.1016/j.tics.2011.12.010

Chang, C., Leopold, D. A., Schölvinck, M. L., Mandelkow, H., Picchioni, D., Liu, X., Ye, F. Q., Turchi, J. N., & Duyn, J. H. (2016). Tracking brain arousal fluctuations with fMRI. Proceedings of the National Academy of Sciences of the United States of America, 113, 4518–4523. 10.1073/pnas.1520613113

Christoff, K., Gordon, A. M., Smallwood, J., Smith, R., & Schooler, J. W. (2009). Experience sampling during fMRI reveals default network and executive system contributions to mind wandering. Proceedings of the National Academy of Sciences of the United States of America, 106, 8719–8724. 10.1073/pnas.0900234106

Corbetta, M., Patel, G., & Shulman, G. L. (2008). The reorienting system of the human brain: From environment to theory of mind. Neuron, 58, 306–324. 10.1016/j.neuron.2008.04.017

de Gee, J. W., Correa, C. M. C., Weaver, M., Donner, T. H., & van Gaal, S. (2021). Pupil dilation and the slow wave ERP reflect surprise about choice outcome resulting from intrinsic variability in decision confidence. Cerebral Cortex, 31(7), 3565–3578. 10.1093/cercor/bhab032

Eickhoff, S. B., Stephan, K. E., Mohlberg, H., Grefkes, C., Fink, G. R., Amunts, K., & Zilles, K. (2005). A new SPM toolbox for combining probabilistic cytoarchitectonic maps and functional imaging data. NeuroImage, 25, 1325–1335. 10.1016/j.neuroimage.2004.12.034

Esteban, O., Markiewicz, C. J., Blair, R. W., Moodie, C. A., Isik, A. I., Erramuzpe, A., Kent, J. D., Goncalves, M., DuPre, E., Snyder, M., Oya, H., Ghosh, S. S., Wright, J., Durnez, J., Poldrack, R. A., & Gorgolewski, K. J. (2019). fMRIPrep: A robust preprocessing pipeline for functional MRI. Nature Methods, 16, 111–116. 10.1038/s41592-018-0235-4

Found, A., & Müller, H. J. (1996). Searching for unknown feature targets on more than one dimension: Investigating a “dimension-weighting” account. Perception & Psychophysics, 58, 88–101. 10.3758/BF03205479

Fox, M. D., Snyder, A. Z., Vincent, J. L., Corbetta, M., Van Essen, D. C., & Raichle, M. E. (2005). The human brain is intrinsically organized into dynamic, anticorrelated functional networks. Proceedings of the National Academy of Sciences of the United States of America, 102, 9673–9678. 10.1073/pnas.0504136102

Gaspelin, N., & Luck, S. J. (2018). The role of inhibition in avoiding distraction by salient stimuli. Trends in Cognitive Sciences, 22, 79–92. 10.1016/j.tics.2017.11.001

Geyer, T., Müller, H. J., & Krummenacher, J. (2008). Expectancies modulate attentional capture by salient color singletons. Vision Research, 48, 1315–1326. 10.1016/j.visres.2008.02.006

Goschy, H., Bakos, S., Müller, H. J., & Zehetleitner, M. (2014). Probability cueing of distractor locations: both intertrial facilitation and statistical learning mediate interference reduction. Frontiers in Psychology, 5, 1195. 10.3389/fpsyg.2014.01195

Gu, Y., Han, F., & Liu, X. (2019). Arousal contributions to resting-state fMRI connectivity and dynamics. Frontiers in Neuroscience, 13, 1190. 10.3389/fnins.2019.01190

Jenkinson, M., Beckmann, C. F., Behrens, T. E. J., Woolrich, M. W., & Smith, S. M. (2012). FSL. NeuroImage, 62, 782–790. 10.1016/j.neuroimage.2011.09.015

Kamp, T., Sorger, B., Benjamins, C., Hausfeld, L., & Goebel, R. (2018). The pre-stimulus default mode network state predicts cognitive task performance levels on a mental rotation task. Brain and Behavior, 8, e01034. 10.1002/brb3.1034

Knapen, T., de Gee, J. W., Brascamp, J., Nuiten, S., Hoppenbrouwers, S. S., & Theeuwes, J. (2016). Cognitive and ocular factors jointly determine pupil responses under equiluminance. PLOS ONE, 11(5), Article e0155574. 10.1371/journal.pone.0155574

Konishi, M., McLaren, D. G., Engen, H., & Smallwood, J. (2015). Shaped by the past: the default mode network supports cognition that is independent of immediate perceptual input. PLOS ONE, 10, e0132209. 10.1371/journal.pone.0132209

Kumada, T. (1999). Limitations in attending to a feature value for overriding stimulus-driven interference. Perception & Psychophysics, 61, 61–79. 10.3758/bf03211949

Landau, A. N., Schreyer, H. M., van Pelt, S., & Fries, P. (2015). Distributed attention is implemented through theta-rhythmic gamma modulation. Current Biology, 25(17), 2332– 2337. 10.1016/j.cub.2015.07.048

Leber, A. B. (2010). Neural predictors of within-subject fluctuations in attentional control. The Journal of Neuroscience, 30, 11458–11465. 10.1523/JNEUROSCI.0809-10.2010

Liesefeld, H. R., Lamy, D., Gaspelin, N., Geng, J. J., Kerzel, D., Schall, J. D., Allen, H. A., Anderson, B. A., Boettcher, S., Busch, N. A., Carlisle, N. B., Colonius, H., Draschkow, D., Egeth, H., Leber, A. B., Müller, H. J., Röer, J. P., Schubö, A., Slagter, H. A., … Wolfe, J. (2024). Terms of debate: Consensus definitions to guide the scientific discourse on visual distraction. Attention, Perception, & Psychophysics, 86, 1445–1472. 10.3758/s13414-023-02820-3

Liesefeld, H. R., Liesefeld, A. M., & Müller, H. J. (2019). Distractor-interference reduction is dimensionally constrained. Visual Cognition, 27, 247–259. 10.1080/13506285.2018.1561568

Liesefeld, H. R., & Müller, H. J. (2019). Distractor handling via dimension weighting. Current Opinion in Psychology, 29, 160–167. 10.1016/j.copsyc.2019.03.003

Liesefeld, H. R., & Müller, H. J. (2021). Modulations of saliency signals at two hierarchical levels of priority computation revealed by spatial statistical distractor learning. Journal of Experimental Psychology: General, 150, 710–728. 10.1037/xge0000970

Liu, X., de Zwart, J. A., Schölvinck, M. L., Chang, C., Ye, F. Q., Leopold, D. A., & Duyn, J. H. (2018). Subcortical evidence for a contribution of arousal to fMRI studies of brain activity. Nature Communications, 9, 395. 10.1038/s41467-017-02815-3

Liu, T. T., Nalci, A., & Falahpour, M. (2017). The global signal in fMRI: nuisance or information? NeuroImage, 150, 213–229. 10.1016/j.neuroimage.2017.02.036

Müller, H. J., Geyer, T., Zehetleitner, M., & Krummenacher, J. (2009). Attentional capture by salient color singleton distractors is modulated by top-down dimensional set. Journal of Experimental Psychology: Human Perception and Performance, 35, 1–16. 10.1037/0096-1523.35.1.1

Müller, H. J., Heller, D., & Ziegler, J. (1995). Visual search for singleton feature targets within and across feature dimensions. Perception & Psychophysics, 57, 1–17. 10.3758/bf03211845

Müller, H. J., Töllner, T., Zehetleitner, M., Geyer, T., Rangelov, D., & Krummenacher, J. (2010). Dimension-based attention modulates feed-forward visual processing. Acta Psychologica, 135, 117–122. 10.1016/j.actpsy.2010.05.004

Murphy, P. R., O’Connell, R. G., O’Sullivan, M., Robertson, I. H., & Balsters, J. H. (2014). Pupil diameter covaries with BOLD activity in human locus coeruleus. Human Brain Mapping, 35, 4140–4154. 10.1002/hbm.22466

Pelli, D. G. (1997). The VideoToolbox software for visual psychophysics: Transforming numbers into movies. Spatial Vision, 10, 437–442. 10.1163/156856897X00366

Pollmann, S., Weidner, R., Müller, H. J., & von Cramon, D. Y. (2000). A fronto-posterior network involved in visual dimension changes. Journal of Cognitive Neuroscience, 12, 480–494. 10.1162/089892900562156

Power, J. D., Lynch, C. J., Dubin, M. J., Silver, B. M., Martin, A., & Jones, R. M. (2020). Characteristics of respiratory measures in young adults scanned at rest, including systematic changes and “missed” deep breaths. NeuroImage, 204, 116234. 10.1016/j.neuroimage.2019.116234

Power, J. D., Plitt, M., Laumann, T. O., & Martin, A. (2017). Sources and implications of whole-brain fMRI signals in humans. NeuroImage, 146, 609–625. 10.1016/j.neuroimage.2016.09.038

Raichle, M. E., MacLeod, A. M., Snyder, A. Z., Powers, W. J., Gusnard, D. A., & Shulman, G. L. (2001). A default mode of brain function. Proceedings of the National Academy of Sciences of the United States of America, 98, 676–682. 10.1073/pnas.98.2.676

Rouder, J. N., Speckman, P. L., Sun, D., Morey, R. D., & Iverson, G. (2009). Bayesian t tests for accepting and rejecting the null hypothesis. Psychonomic Bulletin & Review, 16, 225–237. 10.3758/PBR.16.2.225

Sauter, M., Hanning, N. M., Liesefeld, H. R., & Müller, H. J. (2021). Post-capture processes contribute to statistical learning of distractor locations in visual search. Cortex, 135, 108–126. 10.1016/j.cortex.2020.11.016

Sauter, M., Liesefeld, H. R., & Müller, H. J. (2019). Learning to suppress salient distractors in the target dimension: Region-based inhibition is persistent and transfers to distractors in a nontarget dimension. Journal of Experimental Psychology: Learning, Memory, and Cognition, 45(11), 2080–2097. 10.1037/xlm0000691

Sauter, M., Liesefeld, H. R., Zehetleitner, M., & Müller, H. J. (2018). Region-based shielding of visual search from salient distractors: Target detection is impaired with same-but not different-dimension distractors. Attention, Perception, & Psychophysics, 80, 622–642. 10.3758/s13414-017-1477-4

Schaefer, A., Kong, R., Gordon, E. M., Laumann, T. O., Zuo, X.-N., Holmes, A. J., Eickhoff, S. B., & Yeo, B. T. T. (2018). Local-global parcellation of the human cerebral cortex from intrinsic functional connectivity MRI. Cerebral Cortex, 28, 3095–3114. 10.1093/cercor/bhx179

Schölvinck, M. L., Maier, A., Ye, F. Q., Duyn, J. H., & Leopold, D. A. (2010). Neural basis of global resting-state fMRI activity. Proceedings of the National Academy of Sciences of the United States of America, 107, 10238–10243. 10.1073/pnas.0913110107

Serences, J. T., Shomstein, S., Leber, A. B., Golay, X., Egeth, H. E., & Yantis, S. (2005). Coordination of voluntary and stimulus-driven attentional control in human cortex. Psychological Science, 16, 114–122. 10.1111/j.0956-7976.2005.00791.x

Shi, Z., Zhang, B., Allenmark, F., Yu, H., & Müller, H. J. (2026). Dataset: Retinotopic mapping of statistical learning of distractor suppression in visual search [Data set]. 10.12751/g-node.zfrem9

Spagna, A., Liu, J., & Bartolomeo, P. (2026). Beta and gamma dynamics in attentional networks predict conscious reports. The Journal of Neuroscience, 46(21), Article e0590252026. 10.1523/JNEUROSCI.0590-25.2026

Turchi, J., Chang, C., Ye, F. Q., Russ, B. E., Yu, D. K., Cortes, C. R., Monosov, I. E., Duyn, J. H., & Leopold, D. A. (2018). The basal forebrain regulates global resting-state fMRI fluctuations. Neuron, 97, 940–952.e4. 10.1016/j.neuron.2018.01.032

van Moorselaar, D., & Slagter, H. A. (2019). Learning what is irrelevant or relevant: Expectations facilitate distractor inhibition and target facilitation through distinct neural mechanisms. The Journal of Neuroscience, 39, 6953–6967. 10.1523/JNEUROSCI.0593-19.2019

Wang, B., & Theeuwes, J. (2018). How to inhibit a distractor location? Statistical learning versus active, top-down suppression. Attention, Perception, & Psychophysics, 80, 860–870. 10.3758/s13414-018-1493-z

Wei, P., Yu, H., Müller, H. J., Pollmann, S., & Zhou, X. (2019). Differential brain mechanisms for processing distracting information in task relevant and irrelevant dimensions in visual search. Human brain mapping, 40(1), 110–124. 10.1002/hbm.24358

Weissman, D. H., Roberts, K. C., Visscher, K. M., & Woldorff, M. G. (2006). The neural bases of momentary lapses in attention. Nature Neuroscience, 9, 971–978. 10.1038/nn1727

Wong, C. W., Olafsson, V., Tal, O., & Liu, T. T. (2013). The amplitude of the resting-state fMRI global signal is related to EEG vigilance measures. NeuroImage, 83, 983–990. 10.1016/j.neuroimage.2013.07.057

Yang, X., Fiebelkorn, I. C., Jensen, O., Knight, R. T., & Kastner, S. (2024). Differential neural mechanisms underlie cortical gating of visual spatial attention mediated by alpha-band oscillations. Proceedings of the National Academy of Sciences of the United States of America, 121(45), Article e2313304121. 10.1073/pnas.2313304121

Yellin, D., Berkovich-Ohana, A., & Malach, R. (2015). Coupling between pupil fluctuations and resting-state fMRI uncovers a slow build-up of antagonistic responses in the human cortex. NeuroImage, 106, 414–427. 10.1016/j.neuroimage.2014.11.034

Yeo, B. T. T., Krienen, F. M., Sepulcre, J., Sabuncu, M. R., Lashkari, D., Hollinshead, M., Roffman, J. L., Smoller, J. W., Zöllei, L., Polimeni, J. R., Fischl, B., Liu, H., & Buckner, R. L. (2011). The organization of the human cerebral cortex estimated by intrinsic functional connectivity. Journal of Neurophysiology, 106, 1125–1165. 10.1152/jn.00338.2011

Zehetleitner, M., Goschy, H., & Müller, H. J. (2012). Top-down control of attention: it’s gradual, practice-dependent, and hierarchically organized. Journal of Experimental Psychology: Human Perception and Performance, 38, 941–957. 10.1037/a0027629

Zhang, B., Allenmark, F., Liesefeld, H. R., Shi, Z., & Müller, H. J. (2019). Probability cueing of singleton-distractor locations in visual search: Priority-map-versus dimension-based inhibition? Journal of Experimental Psychology: Human Perception and Performance, 45, 1146–1163. 10.1037/xhp0000652

Zhang, B., Weidner, R., Allenmark, F., Bertleff, S., Fink, G. R., Shi, Z., & Müller, H. J. (2022). Statistical learning of frequent distractor locations in visual search involves regional signal suppression in early visual cortex. Cerebral Cortex, 32, 2729–2744. 10.1093/cercor/bhab377

