## Supplementary for "Pre-stimulus cortical state predicts context-dependent expression of learned distractor suppression"

### **Supplementary Results**

#### **S1. Neural substrates of same-dimension distractor responses**

**Table S1.** Same-dimension > absent activation clusters (C1; *z* > 3.09, *p* < .001).

| **Region** | **Hemisphere** | **Peak z** | **x** | **y** | **z** | **Cluster size**  **(voxels)** |
| --- | --- | --- | --- | --- | --- | --- |
| FEF | L | 4.63 | −37 | −1 | 60 | 124 |
| IPS/SPL | L | 4.54 | −34 | −58 | 50 | 156 |
| SMA | Bilateral | 4.43 | 6 | 21 | 44 | 98 |
| SPL | R | 4.25 | 18 | −67 | 53 | 87 |
| IPL | L | 4.04 | −52 | −40 | 46 | 72 |
| IFG | R | 3.85 | 42 | 8 | 27 | 54 |
| Thalamus | L | 3.72 | −12 | −18 | 8 | 38^a^ |
| Thalamus | R | 3.54 | 14 | −16 | 6 | 37^a^ |

Note. All clusters survived the cluster-level FWE criterion except the thalamic clusters. *N* = 34. Coordinates are reported in MNI space.

#### **S2. Whole-brain location-learning effects**

**Table S2.** Descriptive whole-brain clusters for the location-learning contrast (Rare > Frequent; SS group, N = 17; z > 3.09, *p* < .001).

| **Region** | **Hemisphere** | **Peak z** | **x** | **y** | **z** | **Cluster size**  **(voxels)** |
| --- | --- | --- | --- | --- | --- | --- |
| Occipital cortex | R | 3.95 | 32 | −97 | -9 | 28 |
| Lateral occipital complex | L | 3.71 | −54 | −70 | -9 | 39 |
| Occipital cortex | L | 3.70 | -28 | -97 | -16 | 13 |

#### Note. This SS-group whole-brain map is retained as a descriptive visualization of the visual-cortical location effect.

#### **S3. Exploratory whole-brain group-difference analysis**

To identify regions where the SS and DS groups differed in overall activation, we conducted an exploratory whole-brain second-level analysis of the main effect of group, averaging across all conditions within each participant. No group-difference clusters survived cluster-level FWE correction after applying the voxelwise threshold z > 3.09 (*p* < .001). We therefore report exploratory clusters at *p* < .001 and *k* > 15 for descriptive context (**Table S3**). We then extracted GS modulation from all SS > DS group-difference peaks to test whether regions with overall group differences also showed differential GS modulation. Left medial frontal cortex showed the strongest group difference in GS modulation (*p* = .025, uncorrected; **Figure S1**).

**Table S3.** Exploratory group differences in neural activation.

| **Region** | **Hemisphere** | **Peak z** | **x** | **y** | **z** | **Cluster size**  **(voxels)** |
| --- | --- | --- | --- | --- | --- | --- |
| Postcentral gyrus | R | 4.67 | 38 | −30 | 50 | 26 |
| Medial frontal gyrus (MFG) | L | 3.92 | −18 | 50 | 30 | 15 |
| Superior Frontal Gyrus (SFG) | R | 3.62 | 26 | 36 | 44 | 21 |
| Parahippocampal gyrus | L | 3.50 | -24 | -52 | -6 | 30 |

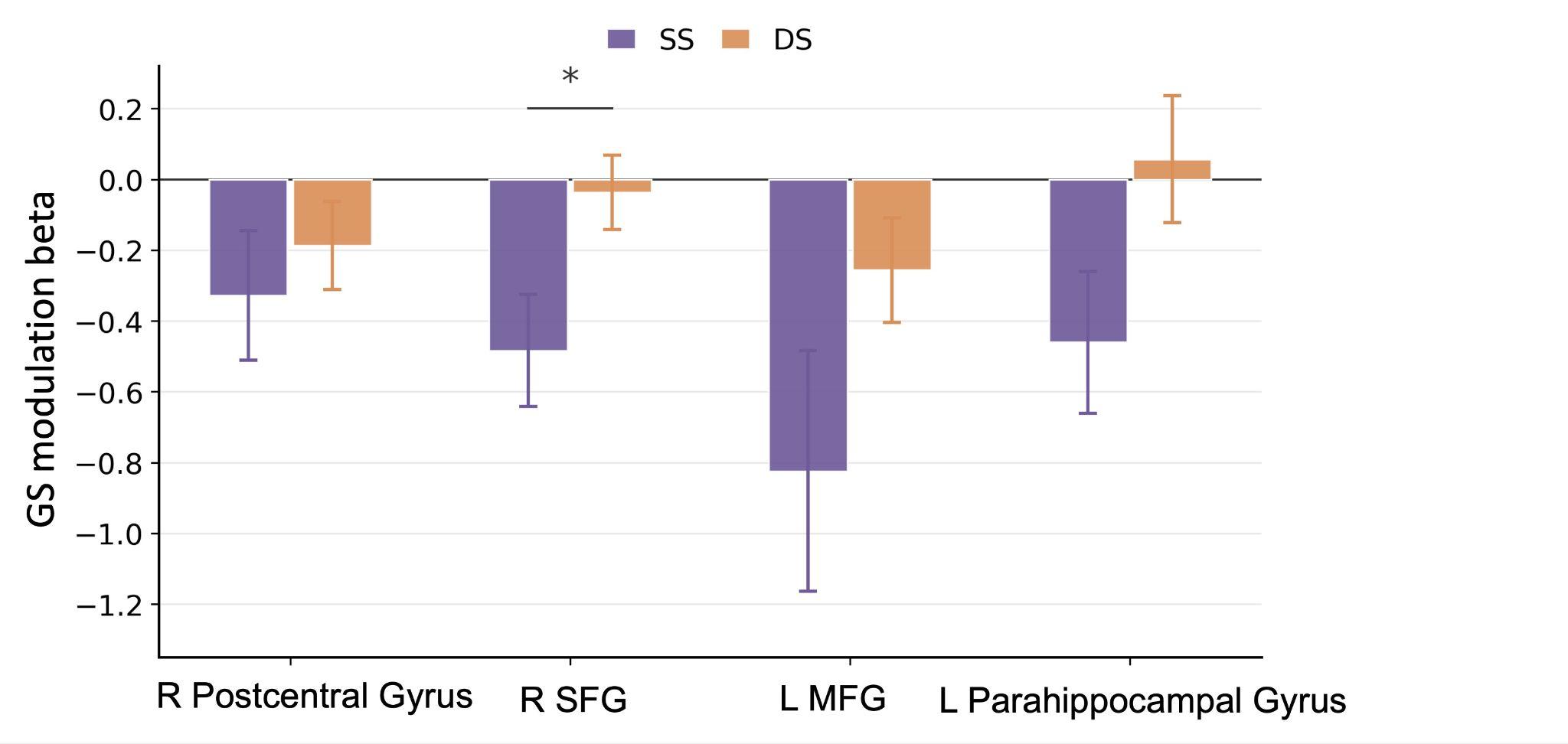

**Figure S1.** GS modulation extracted from group-difference peaks. Effects were numerically stronger in the SS group (orange) than in the DS group (purple), with left medial frontal cortex reaching uncorrected significance (*p* = .025). * *p* < .05, uncorrected.

#### **S4. GS robustness and alternative model checks**

Because GS can carry motion- and physiology-related variance, we tested whether the GS-RT pattern survived nuisance controls. Trial-level pre-stimulus GS covaried significantly with all three nuisance proxies over the same pre-stimulus window: framewise displacement (FD), *β* = .030, *p* < .001; CSF signal, *β* = .393, *p* < .001; and white-matter (WM) signal, *β* = .424, *p* < .001. The behavioral GS effect nevertheless remained stable. The key trial-level GS × Group contrast remained significant when the model controlled for previous trial type alone (*β* = 19.32 ms/*SD*, *p* = .005) and when it additionally controlled for FD, CSF, and WM (*β* = 18.88 ms/*SD*, *p* = .006). The omnibus GS × Group terms likewise remained significant without nuisance proxies, *χ*²(3) = 25.34, *p* < .001, and with them included, *χ*²(3) = 23.74, *p* < .001. Thus, although GS covaried with nuisance proxies, previous trial type, FD, CSF, and WM did not trivially explain the behavioral GS effect. Adding pre-display microsaccade and overt-saccade rates to the RT model did not attenuate the GS × Group association: the coefficient changed from *β* = 25.84 ms, *p* = .001, to *β* = 25.90 ms, *p* = .001, corresponding to −0.2% attenuation. Neither eye-rate × Group interaction survived correction. Accounting for distractor capture at the first saccade reduced the GS × Group RT coefficient by only 4.5%, and the broader first-saccade adjustment reduced it by only 5.5%, leaving the interaction reliable.

The opposite-sign GS-RT pattern also argued against a simple task-difficulty account, which would predict effects in the same direction for both groups, differing mainly in magnitude. Controlling for subject-level distractor-cost magnitude (same-dimension RT minus absent RT) did not eliminate the GS × Group interaction (p < .001). Because this robustness model used a different coefficient parameterization from the primary joint model, we focus on the persistence of the interaction rather than comparing the signed coefficient directly with the primary estimate.

Finally, the GS-RT relationship did not show reliable evidence for an inverted-U state-performance curve. Adding a quadratic GS term (GS²) produced neither a significant main quadratic effect, *β* = +3.0 ms, 95% *CI* [−1.2, 7.2], *SE* = 2.14, *p* = .158, nor a significant GS² × Group interaction, *β* = −3.9 ms, 95% *CI* [−9.9, 2.1], *SE* = 3.08, *p* = .211. These results remain compatible with a monotonic GS-RT relationship in this task.

**S5. Multimodal eye-movement sensitivity analyses**

Fixation was instructed but not gaze-contingently enforced. Excluding trials containing a >2° saccade during the response-bounded display period removed 21.0% trials. The pooled same-dimension capture effect remained large (163.4 ms before exclusion vs. 155.9 ms afterward); the 7.5-ms reduction was not significant, 95% CI [−28.7, 13.8], p = .477. The Same-minus-Different effect was similarly preserved (160.2 vs. 154.4 ms; change = −5.8 ms, 95% CI [−19.4, 7.9], p = .392). Within SS, the Rare-minus-Frequent RT effect remained significant after exclusion (63.3 ms, 95% CI [10.1, 116.6], p = .024), and its 0.7-ms reduction relative to the unfiltered estimate was not significant, p = .908.

For fMRI, we fitted matched unadjusted and saccade-adjusted ROI GLMs in participants with usable scanner gaze. 2,998 trials measured > 2° display-period saccades were modeled as separate HRF-convolved nuisance events at their observed onset. In the primary attention-network ROI, saccade-adjusted estimates remained reliable for C1 (Same − Absent), β = 0.698%, 95% CI [0.267, 1.129], p = .002; C3 (Same − Different), β = 0.666%, 95% CI [0.159, 1.172], p = .010.
